# Neuromechanical strategies for obstacle negotiation during overground locomotion following an incomplete spinal cord injury in adult cats

**DOI:** 10.1101/2023.02.21.529373

**Authors:** Charly G. Lecomte, Stephen Mari, Johannie Audet, Sirine Yassine, Angèle N. Merlet, Caroline Morency, Jonathan Harnie, Claudie Beaulieu, Louis Gendron, Alain Frigon

**Author notes:** **Corresponding author:** Alain Frigon, Ph.D., Université de Sherbrooke, 3001 12e Avenue Nord, Department of Pharmacology-Physiology, Faculty of Medicine and Health Sciences, Sherbrooke, Quebec J1H 5N4, Canada.

## Abstract

Following incomplete spinal cord injury in animals, including humans, substantial locomotor recovery can occur. However, functional aspects of locomotion, such as negotiating an obstacle remains challenging. We collected kinematic and electromyography data in ten adult cats before and at weeks 1-2 and 7-8 after a lateral mid-thoracic hemisection while they negotiated obstacles of three different heights. Intact cats always cleared obstacles without contact. At weeks 1-2 after hemisection, the ipsilesional hindlimb contacted obstacles in ~50% of trials, triggering a stumbling corrective reaction or lack of response. When complete clearance occurred, we observed exaggerated ipsilesional hindlimb flexion when it crossed the obstacle with the contralesional limbs leading. At weeks 7-8 after hemisection, complete clearance increased in favor of absent responses while the proportion of stumbling corrective reactions remained relatively the same. We found redistribution of weight support after hemisection, with reduced diagonal supports and increased homolateral supports, particularly on the contralesional side. The main neural strategy for complete clearance in intact cats consisted of increased activation of muscles that flex the knee. After hemisection, knee flexor activation remained but it was insufficient or more variable as the limb approached the obstacle. Intact cats also increased their speed when stepping over an obstacle, an increase that disappeared after hemisection. The increase in complete clearance over time after hemisection paralleled the recovery of muscle activation patterns or new strategies. Our results suggest partial recovery of anticipatory control through neuroplastic changes in the locomotor control system.

## Introduction

Animals, including humans, require constant adjustments in their gait pattern to navigate in a changing environment. For instance, animals must often step over obstacles, which requires modifying the trajectory of the limb stepping over the obstacle and coordinating the other limbs for balance. Safely negotiating an obstacle involves several levels of the nervous system. When anticipating an obstacle, the visual cortex receives information and sends signals to motor and pre-motor areas [1–5]. In turn, corticospinal and rubrospinal tracts send signals to spinal motor circuits, which then activate muscles and alter limb trajectory to avoid the obstacle [6–9]. If the obstacle is not anticipated, the foot dorsum contacts the obstacle and a reflex mechanism, triggered by cutaneous afferents, modifies limb trajectory to step over the obstacle and prevent stumbling, termed the stumbling corrective reaction (SCR) [10–18]. The SCR occurs in low-thoracic spinal-transected cats, consistent with a spinal mechanism [19,20].

Following incomplete spinal cord injury (SCI) in humans, although substantial walking recovery can occur, notable deficits persist in features of locomotion critical for community ambulation, including obstacle negotiation [21,22]. Even people categorized as AIS D (American Spinal Injury Association Impairment Scale), the highest level of recovery, fail to properly negotiate obstacles [23]. Despite the importance of safely negotiating obstacles after SCI, few studies have investigated it. Drew *et al*., (1996) showed that cortico- and/or rubrospinal tracts were involved in obstacle negotiation during treadmill locomotion by performing dorsolateral spinal lesions at low thoracic levels (T13) in cats. A SCR emerged after the lesion because cats could not anticipate and avoid the obstacle [reviewed in 24]. Another study conducted during overground locomotion after a T10 lateral hemisection in cats showed that the ipsilesional hindlimb contacted the obstacle in ~90% of trials two weeks after SCI [25]. Despite some recovery in the ability to modify limb trajectory over the course of eight weeks before reaching a plateau, limb contact persisted in ~50% of trials, with a marked inability to lift the paw sufficiently to overcome the obstacle [25]. However, in that study, they did not record muscle activity, although they highlighted the importance of doing so in future studies, and only focused on some kinematic adjustments of the ipsilesional hindlimb.

An important aspect to consider when negotiating an obstacle is the limb that crosses the obstacle first (the leading limb) because the biomechanical demands differ. In humans, despite no difference in toe clearance (i.e. the height of the toe over the obstacle) between the leading and trailing legs, vertical movement of the foot is faster for the trailing limb while the horizontal velocity is higher for the leading limb. When the foot lands after crossing the obstacle, vertical and horizontal velocities are similar to a normal step for the trailing limb while vertical velocity is higher for the leading limb [26]. The strategy for the trailing limb allows sufficient time after limb elevation to decelerate the limb for a smooth foot contact [26] while the leading limb strategy prevents slipping at contact [27]. For the leading limb, foot elevation is mainly driven by hip and knee flexion with more hip elevation than for the trailing limb where the knee and ankle contribute to limb flexion [26]. In cats and humans, flexion of the knee is always the main contributor in vertical elevation [28,29].

The purpose of the present study was to characterize neuromechanical strategies employed by adult cats when negotiating obstacles of different heights during overground locomotion before and after a right lateral hemisection at mid-thoracic levels to determine disruptions in voluntary/anticipatory control and neuroplastic changes in spared structures/pathways over time. We did this by recording electromyography (EMG) and kinematic data in the hindlimbs before (intact) and at 1-2 and 7-8 weeks after hemisection. We also separated trials with right (ipsilesional) and left (contralesional) limbs leading. We hypothesize that the loss in commands from the brain impairs activation of muscles that flex the ipsilesional hindlimb, particularly knee flexors, disrupting the ability to clear an obstacle without contact. However, over time, neuroplastic changes allow for some recovery. We also expect different strategies to clear an obstacle based on whether the ipsilesional hindlimb is the leading or trailing limb due to different biomechanical constraints. We present some of our results using an optimal control framework.

## Results

### Different types of obstacle negotiations after lateral hemisection

In the intact state, all cats cleared the obstacles at all three heights without contact, which we term complete clearance (CC), while after hemisection other negotiation types appeared (Fig. 1A). After hemisection, we observed CC, a stumbling corrective reaction (SCR), where the limb contacts the obstacle and evokes a reflex response to move it away and over the obstacle, as well as an absence of response following contact, which we term ‘Other’, as in [25]. In the intact state, CC represented 100% of trials. After hemisection, the proportion of CC significantly decreased at all three obstacle heights, even for obstacles of 1 cm. At 1-2 weeks after hemisection, CC and SCR represented respectively 39.8% and 39.6% of trials while 20.6% were Other. At weeks 1-2, considering only obstacles of 5 and 9 cm, 56.1% were SCR while the proportion of CC and Other were similar at 21.8% and 22.0%, respectively. At weeks 7-8 after hemisection, the amount of CC significantly increased to 62.9% (P < 0.0001) in favor of Other (P < 0.0001) which almost disappeared with only 3.0% of trials. At weeks 7-8, considering only obstacles of 5 and 9 cm, CC represented 46.1% of trials and SCR 47.1%. The proportion of Other remained low at 4.4% of trials (Fig. 1B). At weeks 7-8, Other trials looked much more like “complete clearance with toe drag” [25]. Thus, Other trials will not be discussed further at weeks 7-8. Table 1 summarizes the negotiation type made by each cat at the three time points.

**Table 1.** Type of obstacle negotiation after lateral hemisection. The table shows the number of negotiation types/total number of cycles made by individual cats at both time points after hemisection. The left column indicates cat names. CC: complete clearance; SCR: stumbling corrective reaction; LLL: Left limbs leading; RLL: Right limbs leading.

|  |  | 1-2 weeks after hemisection |  |  |  |  |  | 7-8 weeks after hemisection |  |  |  |  |  |
| --- | --- | --- | --- | --- | --- | --- | --- | --- | --- | --- | --- | --- | --- |
|  |  | CC |  | SCR |  | Other |  | CC |  | SCR |  | Other |  |
|  |  | LLL | RLL | LLL | RLL | LLL | RLL | LLL | RLL | LLL | RLL | LLL | RLL |
| AR | #1 | 8/23 | 6/23 | 9/23 |  |  |  | 14/28 | 14/28 |  |  |  |  |
|  | #5 | 5/21 |  | 7/21 | 6/21 | 3/21 |  | 13/30 | 5/30 | 4/30 | 5/30 |  | 3/30 |
|  | #9 | 6/21 |  | 2/21 | 9/21 | 4/21 |  | 17/31 | 3/31 |  | 8/29 | 3/31 |  |
| CE | #1 | 6/23 | 11/23 |  |  | 3/23 | 3/23 | 10/25 | 15/25 |  |  |  |  |
|  | #5 | 5/27 | 6/27 | 4/27 | 6/27 | 3/27 | 3/27 | 14/24 | 4/24 | 6/24 |  |  |  |
|  | #9 | 3/24 | 3/24 |  | 10/24 | 5/24 | 3/24 | 16/23 |  |  | 4/23 | 3/23 |  |
| GR | #1 | 17/32 | 9/32 |  |  | 3/32 | 3/32 | 23/31 | 8/31 |  |  |  |  |
|  | #5 | 4/21 |  | 9/21 | 3/21 |  | 5/21 | 18/27 |  | 7/27 | 2/27 |  |  |
|  | #9 | 10/28 |  | 4/28 | 14/28 |  |  | 25/32 |  | 4/32 | 3/32 |  |  |
| HO | #1 | 12/22 | 3/22 |  |  | 3/22 | 4/22 | 10/33 | 11/33 |  | 12/33 |  |  |
|  | #5 |  | 3/25 | 13/25 | 3/25 | 3/25 | 3/25 | 7/25 | 3/25 | 9/25 | 3/25 | 3/25 |  |
|  | #9 |  |  | 3/21 | 7/21 | 7/21 | 4/21 | 12/28 | 3/28 | 3/28 | 10/28 |  |  |
| JA | #1 | 10/31 | 11/31 | 3/31 | 3/31 |  | 4/31 | 13/29 | 13/29 | 3/29 |  |  |  |
|  | #5 | 6/27 | 3/27 | 3/27 | 7/27 | 5/27 | 3/27 | 8/32 | 3/32 | 11/32 | 7/32 |  | 3/32 |
|  | #9 | 6/26 |  | 6/26 | 9/26 | 5/26 |  | 13/37 | 3/37 | 10/37 | 8/37 | 3/37 |  |
| KA | #1 | 18/35 | 11/35 |  | 3/35 |  | 3/35 | 19/29 | 6/29 | 4/29 |  |  |  |
|  | #5 | 6/23 |  | 8/23 | 9/23 |  |  | 4/26 |  | 18/26 | 4/26 |  |  |
|  | #9 | 11/31 |  | 10/31 | 6/31 |  | 3/31 | 19/37 |  | 15/37 | 3/37 |  |  |
| KI | #1 | 15/24 | 6/24 |  |  | 3/24 |  | 16/25 | 6/25 |  | 3/25 |  |  |
|  | #5 | 3/29 | 3/29 | 8/29 | 12/29 |  | 3/29 | 5/35 |  | 13/35 | 17/35 |  |  |
|  | #9 | 8/31 |  | 11/31 | 10/31 |  | 2/31 | 12/23 |  | 3/23 | 8/23 |  |  |
| MB | #1 | 4/24 | 14/24 | 3/24 |  |  | 3/24 | 25/40 | 12/40 |  |  |  | 3/40 |
|  | #5 |  | 5/23 | 5/23 | 7/23 | 3/23 | 3/23 | 6/40 |  | 15/40 | 16/40 | 3/40 |  |
|  | #9 |  |  |  | 14/21 |  | 7/21 | 9/37 | 3/37 | 7/37 | 15/37 | 3/37 |  |
| PO | #1 | 3/25 | 8/25 |  |  | 10/25 | 4/25 | 8/28 | 20/28 |  |  |  |  |
|  | #5 | 8/21 | 4/21 |  |  | 6/21 | 3/21 | 10/29 | 10/29 | 3/29 | 3/29 |  | 3/29 |
|  | #9 | 10/22 |  | 3/22 | 3/22 | 3/22 | 3/22 | 5/25 | 6/25 |  | 5/25 |  | 9/25 |
| UM | #1 | 15/33 | 12/33 | 3/33 |  |  | 3/33 | 17/34 | 17/34 |  |  |  |  |
|  | #5 |  | 6/30 | 14/30 | 7/30 |  | 3/30 | 4/33 | 9/33 | 17/33 | 3/33 |  |  |
|  | #9 | 3/29 |  | 17/29 | 3/29 | 3/29 | 3/29 | 7/27 |  | 12/27 | 8/27 |  |  |

**Figure 1.**
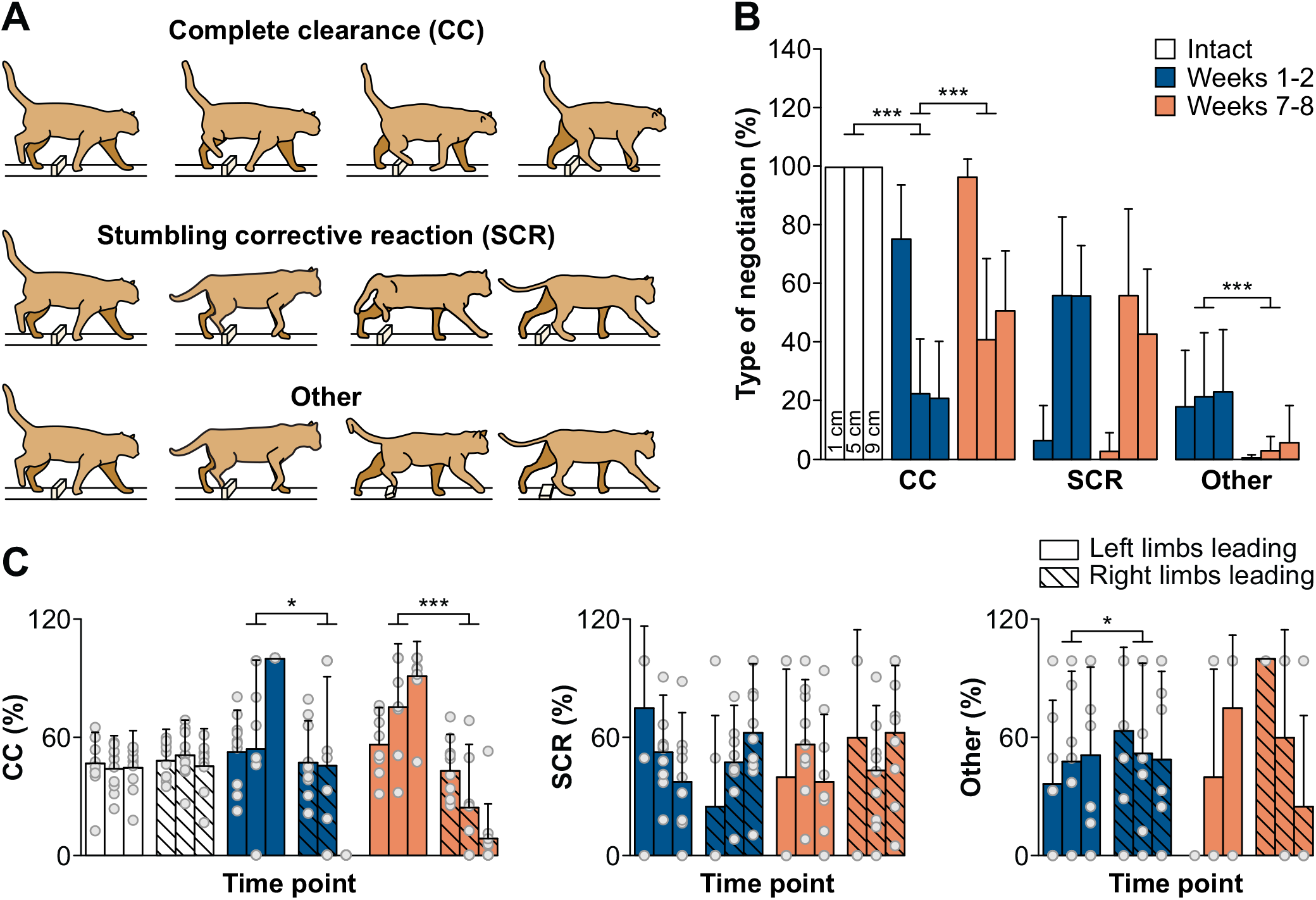
Different types of obstacle negotiations and their proportions. **A**: Schematic illustration of the three obstacle negotiations. **B**: Proportion of each negotiation type for the ipsilesional hindlimb. **C**: Percentage of each negotiation type according to leading limbs. The bars represent mean ± SD for the group while grey circles represent individual data points (mean for each cat). For each obstacle height, we averaged between 3 and 25 trials per cat (n = 10 cats; 5 females and 5 males). When we found a main effect (mixed-effects ANOVA), we performed pairwise comparisons. Asterisks indicate significant difference between sides. * P < 0.01; ** P < 0.001; *** P<0.0001.

For CC in intact cats, we observed no clear side preference, with both Left and Right limbs leading showing similar proportions (Fig. 1C). However, after hemisection, the proportion of CC was significantly greater with Left limbs leading (+102.7% and +202.0% compared to Right limbs leading, respectively, at weeks 1-2 and 7-8). Importantly, no cat performed CC with Right limbs leading for the 9 cm obstacle height at weeks 1-2. SCR at weeks 1-2 and 7-8 after hemisection did not show a side preference in limbs leading. Other at weeks 1-2 after hemisection showed a greater proportion with Right limbs leading (+17.4% compared to Left limbs leading). Furthermore, at weeks 7-8, we observed no Other trials with Left limbs leading at 1 cm (Fig. 1C). After hemisection, the contralesional hindlimb never contacted the obstacle. For all negotiations pooled, we observed no side preference in the intact state (P = 0.3413) and at weeks 1-2 after hemisection (P = 0.2032). However, 7-8 weeks after hemisection, 62.8% of negotiations were with Left limbs leading, which was significantly greater compared to the proportion of Right limbs leading (P = 0.0006).

### Altered and less optimal limb trajectories when negotiating obstacles after hemisection

After hemisection, right hindlimb trajectory was altered compared to the intact state and depended on the type of negotiation, as shown in Figure 2 for a single cat crossing a 5 cm obstacle with Left and Right limbs leading before and at weeks one and seven after hemisection. In the intact state, with Right limbs leading, the right hindlimb begins its swing phase further away from the obstacle compared to Left limbs leading. These distances from the obstacle at swing onset were maintained after hemisection. When a SCR occurred after hemisection, we observed that the right hindpaw was closer to the obstacle at liftoff with Left limbs leading and not sufficiently lifted during swing with Right limbs leading. An absence of a SCR following contact with the obstacle characterized the Other type of negotiation. This is most evident with Right limbs leading at week 1 after hemisection where limb trajectory continued unimpeded following contact and the obstacle fell over.

**Figure 2.**
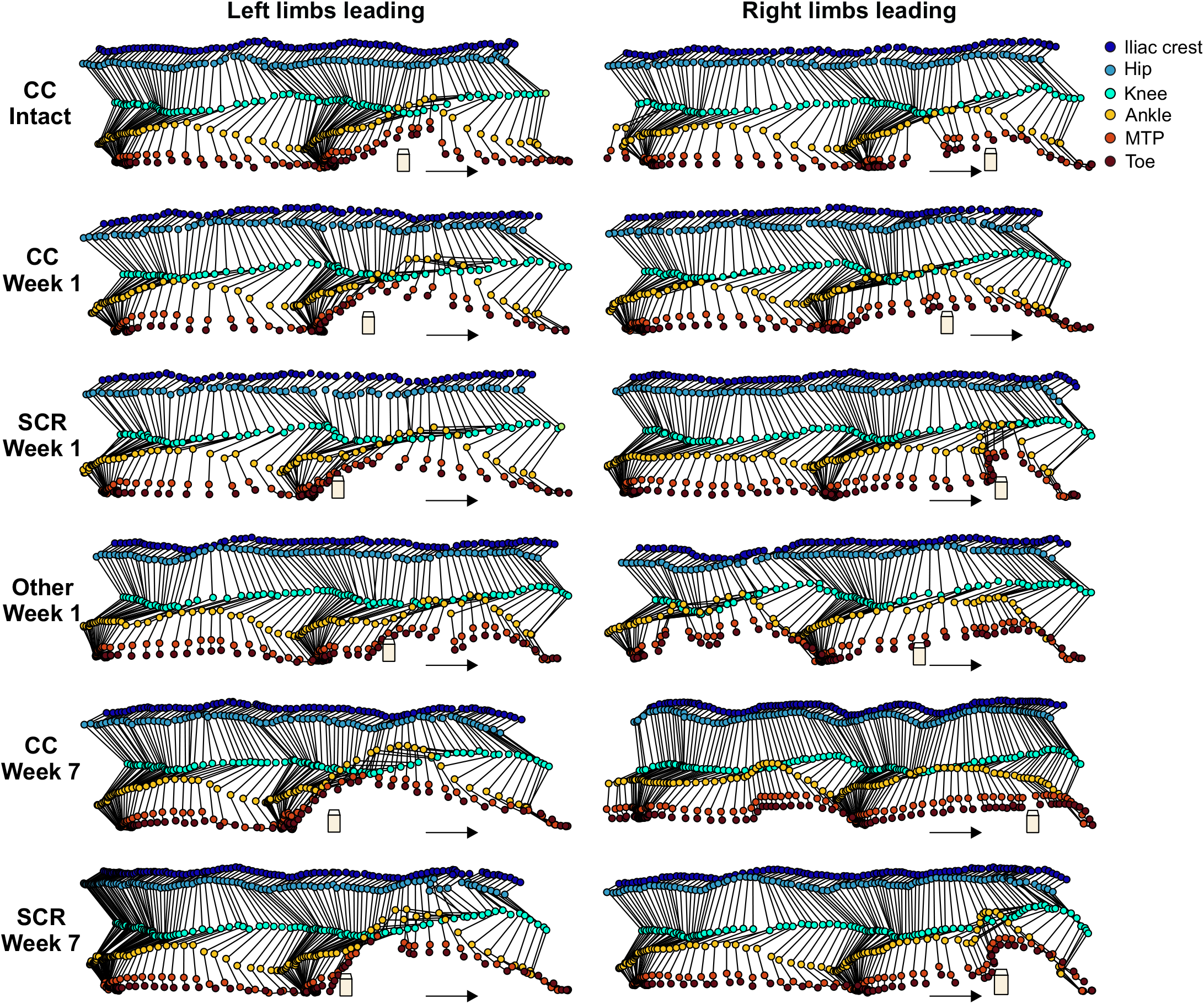
Kinematics of the right hindlimb during obstacle negotiation before and after a spinal hemisection. The figure shows stick figure diagrams of the right hindlimb for representative cycles for the step before and the obstacle step for the six types of negotiation before and at weeks 1 and 7 after hemisection during Left and Right limbs leading. All figures are from cat JA with a 5 cm obstacle. Arrows indicate direction of movement.

When animals step over an obstacle, they must consider various factors, such as the distance between the hindpaw and the obstacle at liftoff (approach distance) and where it lands after clearing it (reception distance). With Left limbs leading (Fig. 3A, left panel), right hindlimb approach distance was shorter for CC at weeks 1-2 and 7-8 weeks after hemisection (P = 0.0011, -43.7% and P < 0.0001, -47.6%, respectively) compared to the intact state, indicating that the right hindpaw was closer to the obstacle. Reception distance was not different after hemisection compared to the intact state, except for SCR at weeks 1-2 with the right hindlimb landing closer to the obstacle (P = 0.0013, -11.8%). With Right limbs leading (Fig 3A, right panel), we found no significant differences for approach distance. The only significant difference was a greater reception distance for SCR at weeks 7-8 after hemisection (P = 0.0007, +24.4%), indicating that the right hindpaw landed further from the obstacle.

**Figure 3.**
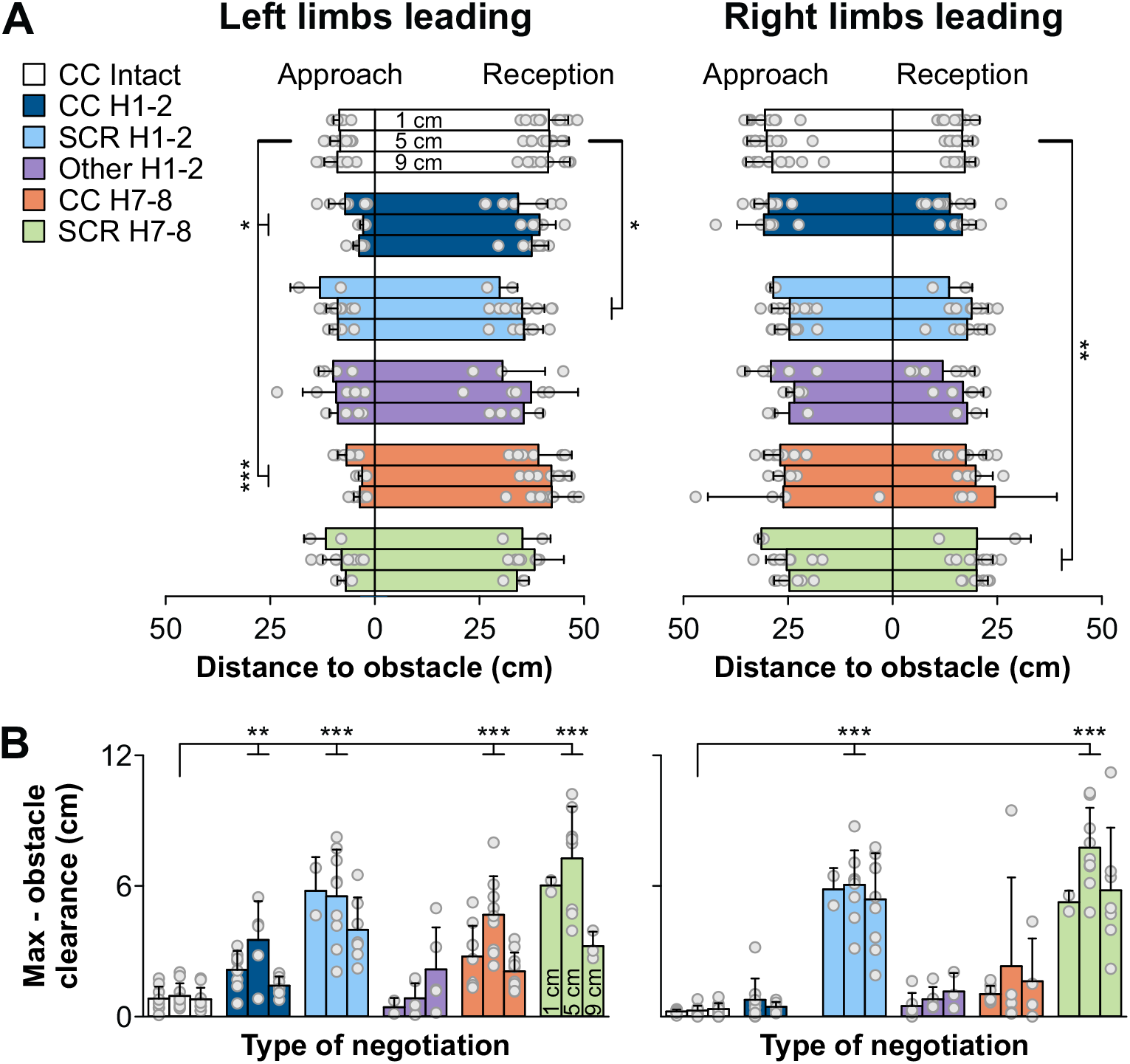
Distances in relation to the obstacle at liftoff and contact and limb elevations before and after a spinal hemisection. **A:** Approach and reception distances of the right hindlimb for each negotiation type before and at weeks 1-2 (H1-2) and 7-8 (H7-8) after hemisection. **B:** Difference between maximal and obstacle clearance of the right hindlimb for each negotiation type. The bars represent mean ± SD for the group while grey circles represent individual data points (mean for each cat). For each obstacle height, we averaged between 3 and 25 trials per cat (n = 10 cats; 5 females and 5 males). When we found a main effect (mixed-effects ANOVA), we performed pairwise comparisons. Asterisks indicate significant difference between complete clearance (CC) in intact cats and the other negotiation types observed after hemisection. SCR, stumbling corrective reaction. * P < 0.01; ** P < 0.001; *** P<0.0001.

Cats and humans will step over an obstacle close to it without contact and optimize maximal flexion to minimize energy expenditure [25]. To assess this and how it might be impaired after hemisection, we measured the difference between the maximal clearance and obstacle clearance for the right hindlimb for each type of negotiation and compared it to CC in the intact state (Fig. 3B). Smaller values reflect a more efficient or optimal obstacle crossing. With Left limbs leading, the difference between maximal and obstacle clearance increased after hemisection for all negotiations except Other at weeks 1-2 (CC: P = 0.0002, +184.6% and SCR: P < 0.0001, +530.9%) and weeks 7-8 (CC: P < 0.0001, +298.7% and SCR: P <0.0001, +686.7%). With Right limbs leading, we observed an increase for SCR at weeks 1-2 and 7-8 (P < 0.0001, +1918.4% and P < 0.0001, +2283.3% respectively) compared to CC intact.

### Negotiating obstacles mainly involves a knee and ankle strategy before and after hemisection

To determine the different strategies used to negotiate obstacles after hemisection, we measured joint angles of the right hindlimb throughout the cycle, as shown for a single cat before and at weeks 1 and 7 after hemisection at an obstacle height of 5 cm (Fig. 4). We compared the step before and when crossing the obstacle. Smaller angle values correspond to greater flexion. In the intact state for this cat, with Left and Right limbs leading, we observed greater angular excursion at the knee and ankle, particularly flexion, in the obstacle step while hip joint excursions were similar. At week 1 after hemisection with Left limbs leading, we observed smaller hip excursion but similar knee and ankle excursions for CC in the obstacle step. With Right limbs leading, all three joints showed greater excursions for CC at week 1 after hemisection in the obstacle step. The SCR at week 1 after hemisection was mainly characterized by greater knee joint excursion in the obstacle step with Left and Right limbs leading. The Other type of negotiation shows little modulation in excursion for all three joints in the obstacle step for Left limbs leading but considerably greater excursions at the hip, knee and ankle with Right limbs leading. At week 7, CC and SCR show increased excursions at all three joints in the obstacle step for Left limbs leading, particularly at the knee and ankle. With Right limbs leading at week 7, CC and SCR were characterized by a greater excursion for the knee and ankle in the obstacle step but not at the hip.

**Figure 4.**
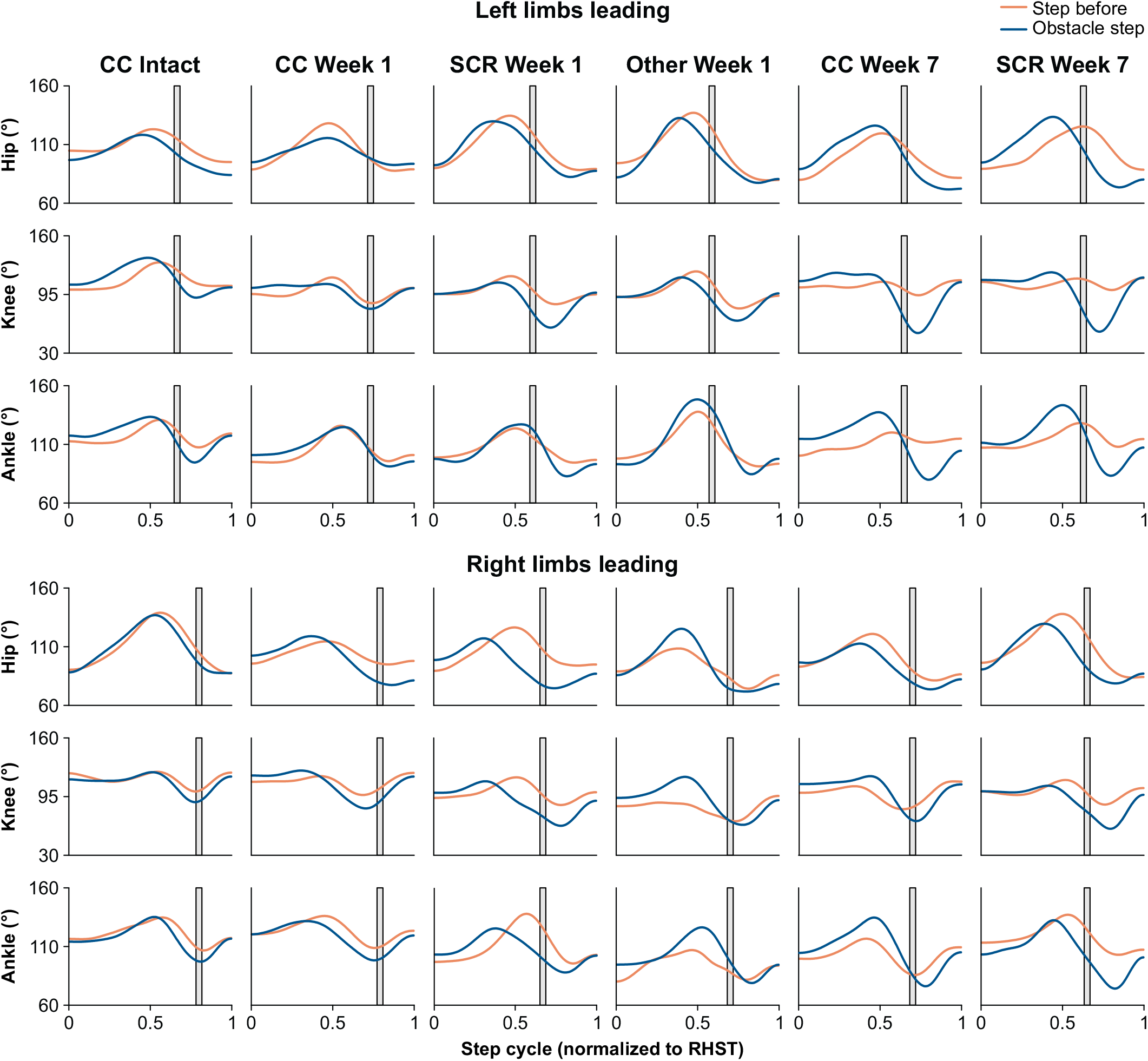
Joint angular excursion of the right hindlimb during obstacle negotiation before and after hemisection in a single cat. Joint angles of the right hip, knee and ankle for the step before and the obstacle step timed to right hindlimb contact and normalized by interpolating the step cycle into 512 equal bins. Each waveform is the average of 3-5 trials from cat JA with a 5 cm obstacle. The gray area represents the timing of the obstacle.

For the group (Fig. 5), hip excursion was not significantly different between the control and obstacle step before or after hemisection in Left and Right limbs leading except for CC at weeks 1-2 (+22.6%) in Right limbs leading. For the knee joint, angular excursion was significantly greater in the obstacle step compared to the control step for all negotiation types before and at weeks 1-2 and 7-8 after hemisection in Left and Right limbs leading. For Left limbs leading, we observed 67.3%, 44.9%, 50.0%, 40.9%, 57.9% and 76.3% increases for intact CC, CC at weeks 1-2, SCR at weeks 1-2, Other at weeks 1-2, CC at weeks 7-8 and SCR at weeks 7-8, respectively. For Right limbs leading, we observed 25.1%, 12.7%, 41.5%, 12.7%, 17.1% and 57.5% increases for intact CC, CC at weeks 1-2, SCR at weeks 1-2, Other at weeks 1-2, CC at weeks 7-8 and SCR at weeks 7-8, respectively. For the ankle joint, angular excursion was significantly greater in the obstacle step compared to the control step for all negotiation types before and at weeks 1-2 and 7-8 after hemisection in Left limbs leading. We observed 23.3%, 16.1%, 29.6%, 26.6%, 27.9% and 36.0% increases for intact CC, CC at weeks 1-2, SCR at weeks 1-2, Other at weeks 1-2, CC at weeks 7-8 and SCR at weeks 7-8, respectively. In Right limbs leading, however, angular excursion was significantly greater in the obstacle step compared to the control step for CC in the intact state (35.9%) and SCR at weeks 1-2 (23.0%) and 7-8 (33.0%) after hemisection.

**Figure 5.**
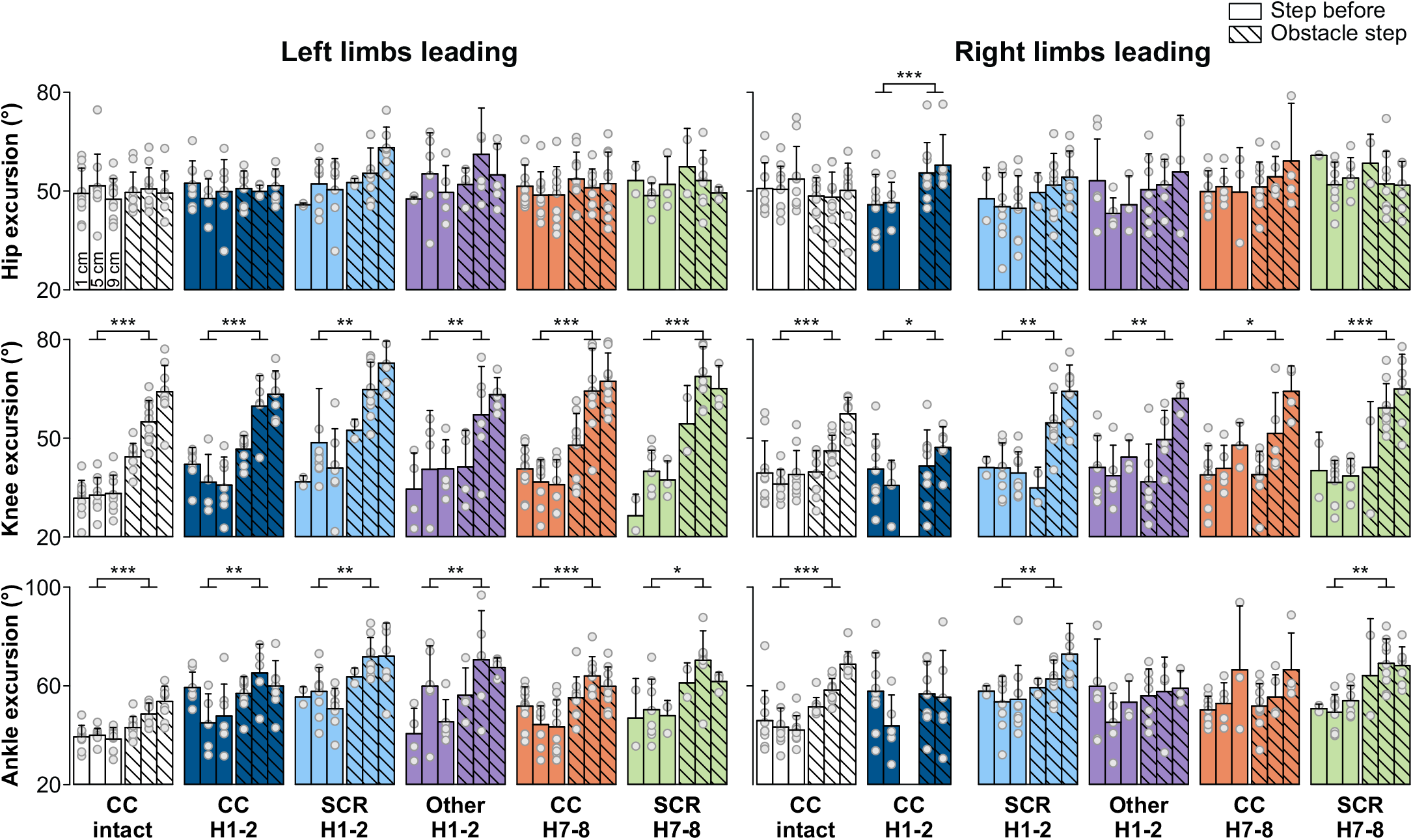
Joint angular excursions of the right hindlimb during obstacle negotiation before and after hemisection for the group. Joint excursions of the right hip, knee and ankle for the step before and the obstacle step for each negotiation type before and at weeks 1-2 (H1-2) and 7-8 (H7-8) after hemisection. The bars represent mean ± SD for the group while grey circles represent individual data points (mean for each cat). For each obstacle height, we averaged between 3 and 25 trials per cat (n = 10 cats; 5 females and 5 males). When we found a main effect (mixed-effects ANOVA), we performed pairwise comparisons. Asterisks indicate significant difference between the step before and the obstacle step. SCR, stumbling corrective reaction. * P < 0.01; ** P < 0.001; *** P < 0.0001.

### The timing of joint flexion depends on the leading limbs after hemisection

To determine if the timing of hip, knee and ankle joint flexion could explain the different negotiation types after hemisection, we measured their flexion onsets in the different negotiations and compared them to CC in the intact state (Fig. 6). In general, for all negotiation types with Left and Right Limbs leading, the hip flexed earlier after hemisection, with the exception of Other at weeks 1-2. With Left limbs leading at weeks 1-2 and 7-8 after hemisection, the hip flexed earlier for CC (P = 0.0053, -5.9% and P < 0.0001, -6.0% respectively) and SCR (P < 0.0001, -9.1% and P < 0.0001, -5.9% respectively). With Right limbs leading at weeks 1-2 and weeks 7-8 after hemisection, the hip flexed earlier for CC (P = 0.0005, -9.5% and P < 0.0001, -10.5%, respectively), SCR (P < 0.0001, -14.4% and P < 0.0001, -13.0%, respectively) and Other at weeks 1-2 (P = 0.0015, -9.0%). Knee flexion onset did not change significantly after hemisection with Left limbs leading and the ankle joint only flexed significantly earlier for CC at weeks 7-8 (P = 0.0019, -3.2%). However, for Right limbs leading, knee flexion occurred earlier for all negotiation types at weeks 1-2 and 7-8 after hemisection, including CC (P= 0.0004, -9.4% and P = 0.0001, -7.9%, respectively), SCR (P < 0.0001, -10.3% and P = 0.0019, -8.4%, respectively) and Other at weeks 1-2 (P = 0.0002, -9.5%). With Right limbs leading, ankle flexion also occurred earlier for CC at weeks 1-2 (P = 0.0005, -6.3%) and SCR at weeks 1-2 and 7-8 (P < 0.0001; -9.8% and P < 0.0001, -7.9%, respectively).

**Figure 6.**
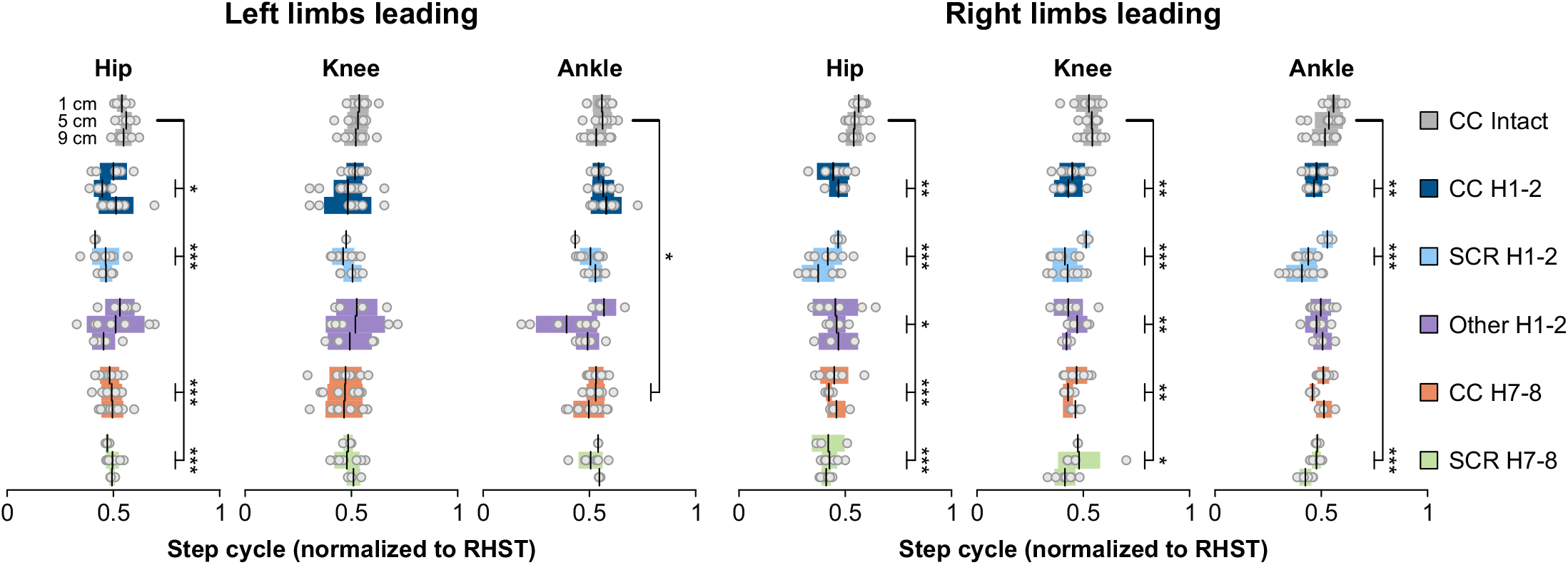
Timing of joint flexions of the right hindlimb during obstacle negotiation before and after hemisection for the group. Timing of flexion onsets of the right hip, knee and ankle for each obstacle height and negotiation type before and at weeks 1-2 (H1-2) and 7-8 (H7-8) after hemisection. The bars represent mean ± SD for the group while grey circles represent individual data points (mean for each cat). For each obstacle height, we averaged between 3 and 25 trials per cat (n = 10 cats; 5 females and 5 males). When we found a main effect (mixed-effects ANOVA), we performed pairwise comparisons. Asterisks indicate significant difference between complete clearance (CC) in intact cats and the other negotiation types observed after hemisection. SCR, stumbling corrective reaction. * P < 0.01; ** P < 0.001; *** P < 0.0001.

### Reorganization of support periods after hemisection

To determine how the four limbs contribute to dynamic balance when stepping over an obstacle and how this is affected by incomplete SCI, we measured support periods before and after hemisection for each negotiation type and compared them to CC in the intact state (Fig. 7). With Left and Right limbs leading, the triple support period involving the two hindlimbs and the left forelimb (Period 1) decreased for all negotiation types after hemisection compared to CC in the intact state (mean -60.0% and -61.2% for Left and Right limbs leading, respectively). We also observed a decrease in both diagonal support periods (Periods 2 and 6) after hemisection for all negotiation types compared to CC in the intact state for Left (mean -55.9% and -61.5% for Periods 2 and 6, respectively) and Right (mean -68.8% and -64.8% for Periods 2 and 6, respectively) limbs leading, with the exception of Other for period 2. In contrast, both homolateral support periods (Periods 4 and 8) increased after hemisection compared to CC in the intact state for Left limbs leading (mean +42.3% and +115.9% for Periods 4 and 8, respectively) with the exception of SCR and Other at 1-2 weeks after hemisection. For Right limbs leading, left homolateral support (Period 8) increased for all negotiation types compared to CC in the intact state (mean +131.6%) while we observed increases for right homolateral support (Period 4) only for CC and SCR at 7-8 weeks after hemisection (mean +40.8%). For Left limbs leading, we also observed a decrease in triple support involving the two forelimbs and the right hindlimb (Period 3) compared to CC in the intact state for CC (−26.8%) and SCR (−30.6%) at weeks 7-8 after hemisection. This was accompanied with an increase in triple support involving the two hindlimbs and the right forelimb (Period 5) for CC (+16.3%) and SCR (+40.8%) at weeks 7-8 after hemisection. For Right limbs leading, we observed an increase in quadrupedal support for SCR at weeks 1-2 (+1208.2%) as well as CC (+1190.3%) and SCR (+1425.4%) at weeks 7-8 after hemisection. Thus, support periods are reorganized after hemisection depending on the type of negotiation and the leading limb.

**Figure 7.**
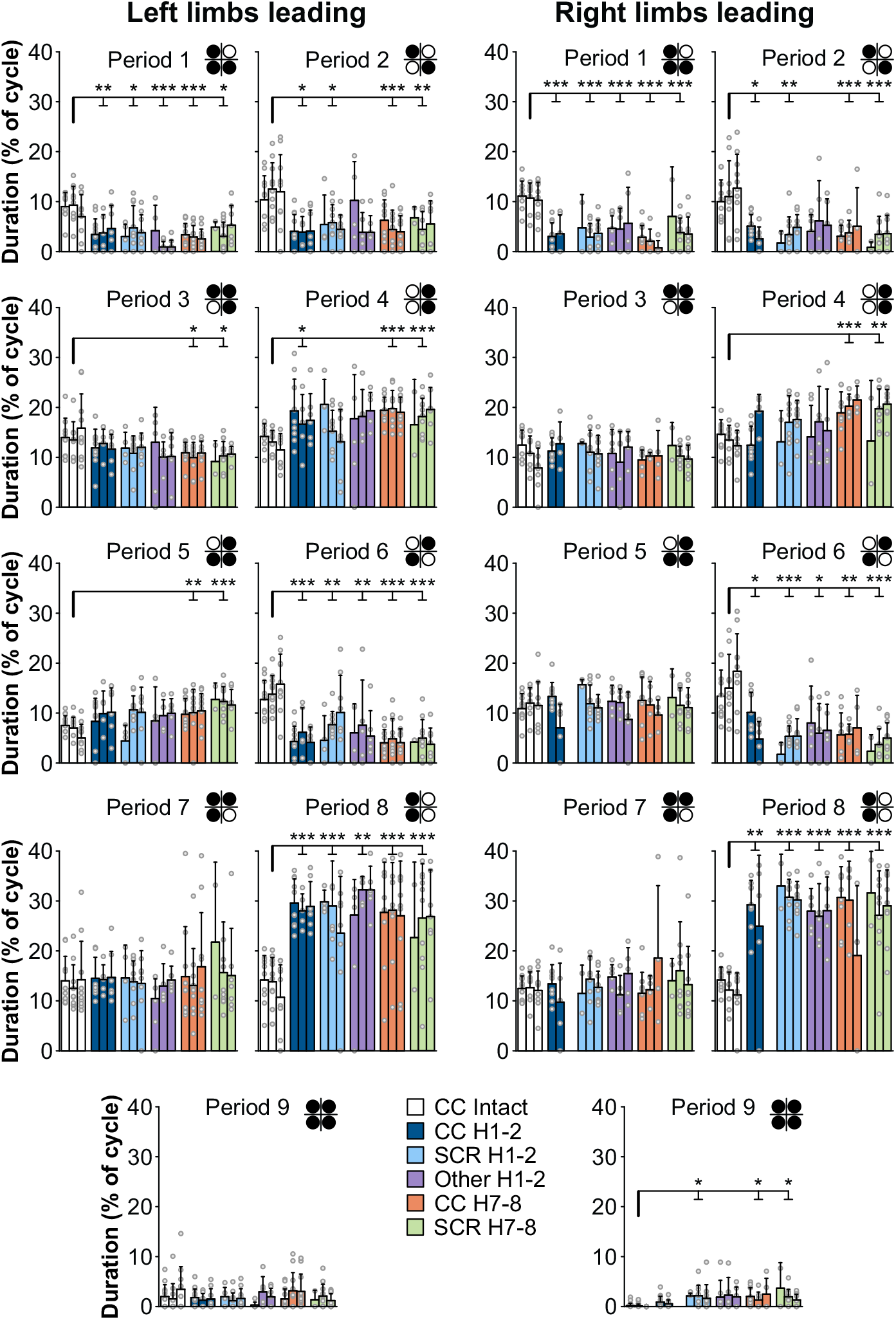
Support periods during obstacle negotiation before and after hemisection for the group. Proportion of each support period normalized to cycle duration for each obstacle height and negotiation type for the obstacle cycle before and at weeks 1-2 (H1-2) and 7-8 (H7-8) after hemisection. The limbs contacting the surface are shown in black in the footfall diagram in each panel. Top left, top right, bottom left and bottom right circles represent left forelimb, right forelimb, left hindlimb and right hindlimb, respectively. The bars represent mean ± SD for the group while grey circles represent individual data points (mean for each cat). For each height, we averaged 3-25 cycles per cat (n = 10 cats; 5 females and 5 males). When we found a main effect (mixed-effects ANOVA), we performed pairwise comparisons. Asterisks indicate significant difference between complete clearance (CC) in intact cats and the other negotiation types observed after hemisection. SCR, stumbling corrective reaction. * P < 0.01; ** P < 0.001; *** P < 0.0001.

### Strategies for stepping over an obstacle involve increasing body speed and stride length

To determine some of the strategies used by cats to step over an obstacle, we measured body speed and stride length during the control and obstacle cycles before and after hemisection. Intact cats performing CC significantly increased their speed when stepping over the obstacle (+7.6% and +22.8% for Left and Right limbs leading, respectively) (Fig. 8A). In contrast, cats did not increase their speed after hemisection in the different negotiation types. Intact cats performing CC significantly increased their stride length when stepping over obstacles (+19.8% and +8.3% for Left and Right Limbs leading, respectively) (Fig. 8B). At weeks 1-2 after hemisection, stride length did not change significantly when stepping over the obstacle for CC, SCR and Other negotiation types in Left limbs leading. We observed an increase in stride length at weeks 7-8 for CC (+17.0%). In Right limbs leading, stride length increased when stepping over the obstacle for all negotiation types at weeks 1-2 after hemisection (CC: +24.5%; SCR: +18.6% and Other: +15.5%). At weeks 7-8, stride length increased only for SCR (+11.3%).

**Figure 8.**
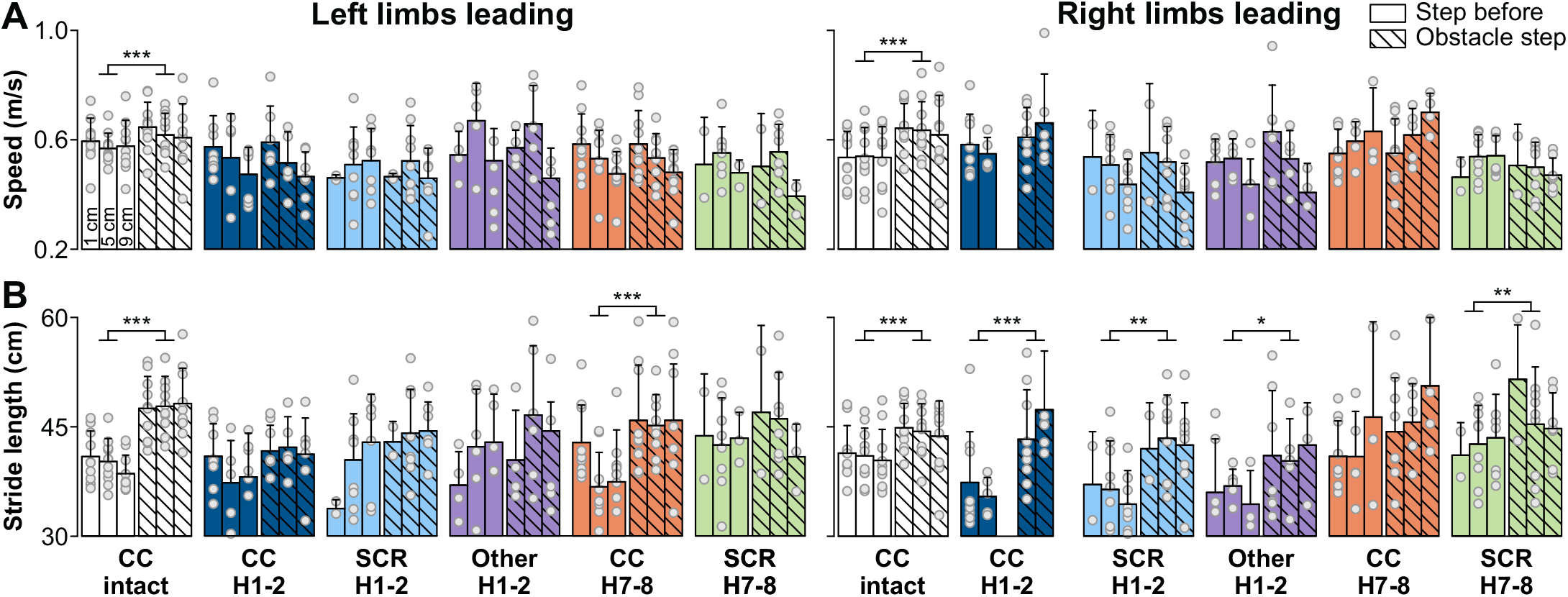
Ipsilesional spatial parameters during obstacle negotiation before and after hemisection. **A:** Body speed and **B:** right hindlimb stride length for the step before and the obstacle step for each obstacle height and negotiation type before and at weeks 1-2 (H1-2) and 7-8 (H7-8) after hemisection. The bars represent mean ± SD for the group while grey circles represent individual data points (mean for each cat). For each height, we averaged 3-25 cycles per cat (n = 10 cats; 5 females and 5 males). When we found a main effect (mixed-effects ANOVA), we performed pairwise comparisons. Asterisks indicate significant difference between the step before and the obstacle step. * P < 0.01; ** P < 0.001; *** P < 0.0001. In the left panel, left limbs are leading and in the right panel right limbs are leading.

### Altered muscle activation strategies when negotiating an obstacle after hemisection

To determine how the nervous system adjusts to a lateral hemisection when negotiating obstacles, we analyzed EMG data for each cat separately to assess individualized strategies and group tendencies. We compared the cycle before (control cycle) and when stepping over the obstacle (obstacle cycle). The muscle activation strategies for Left and Right limbs leading differed to allow the right hindlimb to step over the obstacle in intact cats. Figure 9 shows raw EMG waveforms of four hindlimb muscles bilaterally with the stance phases of the four limbs in one cat for each type of negotiation before and at weeks 1-2 after hemisection at an obstacle height of 5 cm. With Left limbs leading, we observed a large increase in the RST burst and a delay in the activation of the RSRT along with a small increase in its amplitude, allowing the knee to flex and the hip to extend before flexing the hip. Less visible is an increase in RVL amplitude before the right hindlimb steps over the obstacle. With Right limbs leading, we see an increase in RST amplitude combined with concurrent activation of the RSRT and an increase in its amplitude. For complete clearance at weeks 1-2 after hemisection, the RST burst was longer and more variable in both Left and Right limbs leading. The onset of RSRT was also more variable and we did not observe and increase in its amplitude. As a potential compensatory mechanism, we found a second burst in the LSRT during left hindlimb stance in Left limbs leading. This co-activation of hip flexor muscles with other extensors can increase stiffness and limb support. For SCR at weeks 1-2 for Left and Right limbs leading, we see an activation of RST before the obstacle, but it was small compared to the intact state. However, following contact, we observed a large spike in amplitude. The onset of RSRT relative to RST is variable. For the Other type of negotiation, we also observe activation of RST before the obstacle in both conditions but when the right hindpaw contacts the obstacle we see no spike in its amplitude, consistent with a lack of reflex response.

**Figure 9.**
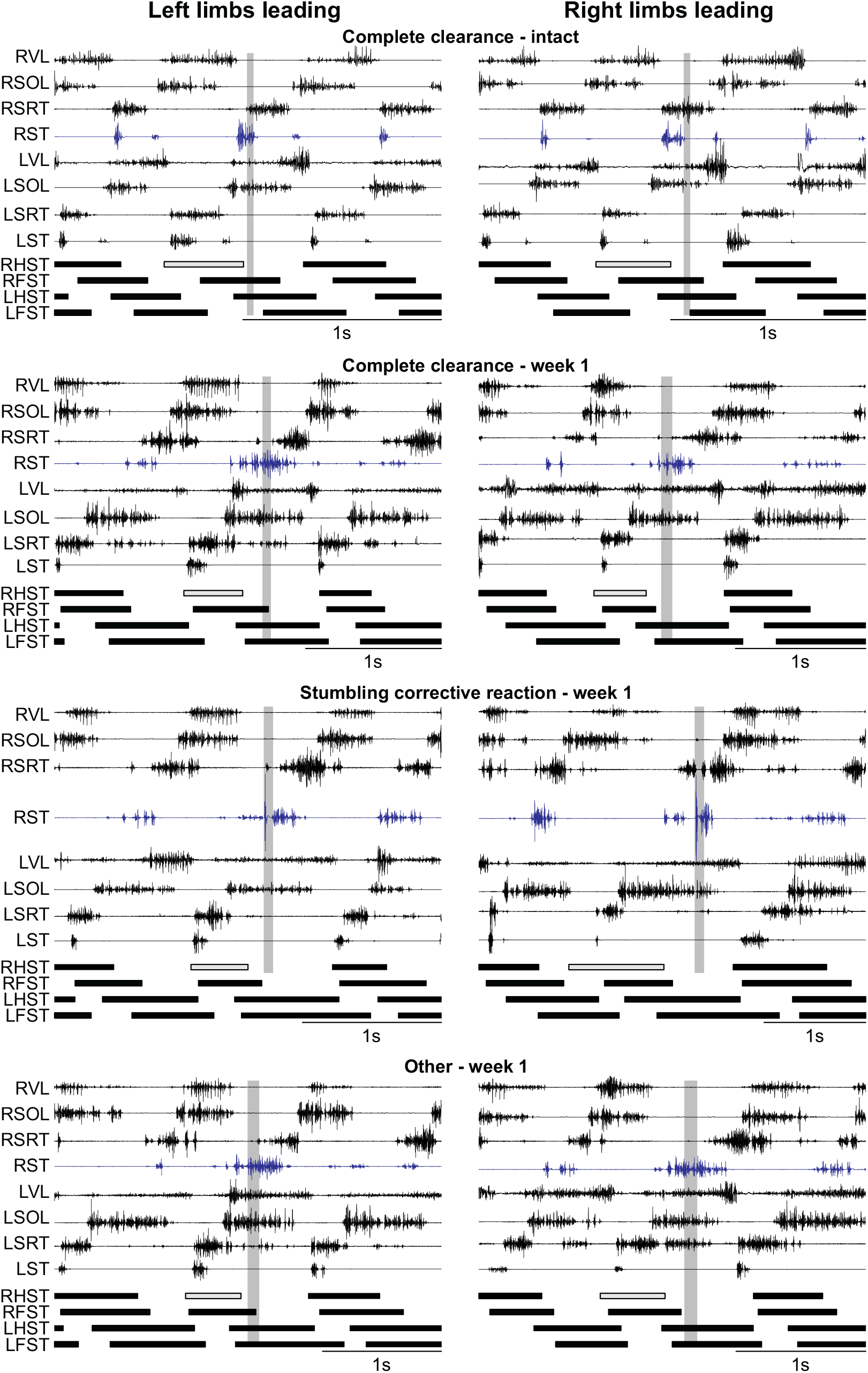
Muscle activity during obstacle negotiation before and after hemisection. Each panel shows the electromyography (EMG) of hindlimb muscles recorded bilaterally (L, left; R, right) along with the stance phases of the four limbs during obstacle negotiation before and at week 1 after hemisection in cat AR at an obstacle height of 5 cm. The gray stance phase represents right hindlimb stance before the obstacle. The vertical scale for each EMG waveform is optimized for viewing. CC, complete clearance; VL, vastus lateralis; SOL, soleus; SRT, anterior sartorius; ST, semitendinosus.

For the group, muscle activation strategies (onsets and amplitudes) emerged from individual cat data, as summarized in Tables 2 and 3 for eight selected muscles before and after hemisection in Left and Right limbs leading, respectively. Remaining muscles can be found in supplementary data (S1 and S2 tables).

**Table 2.**
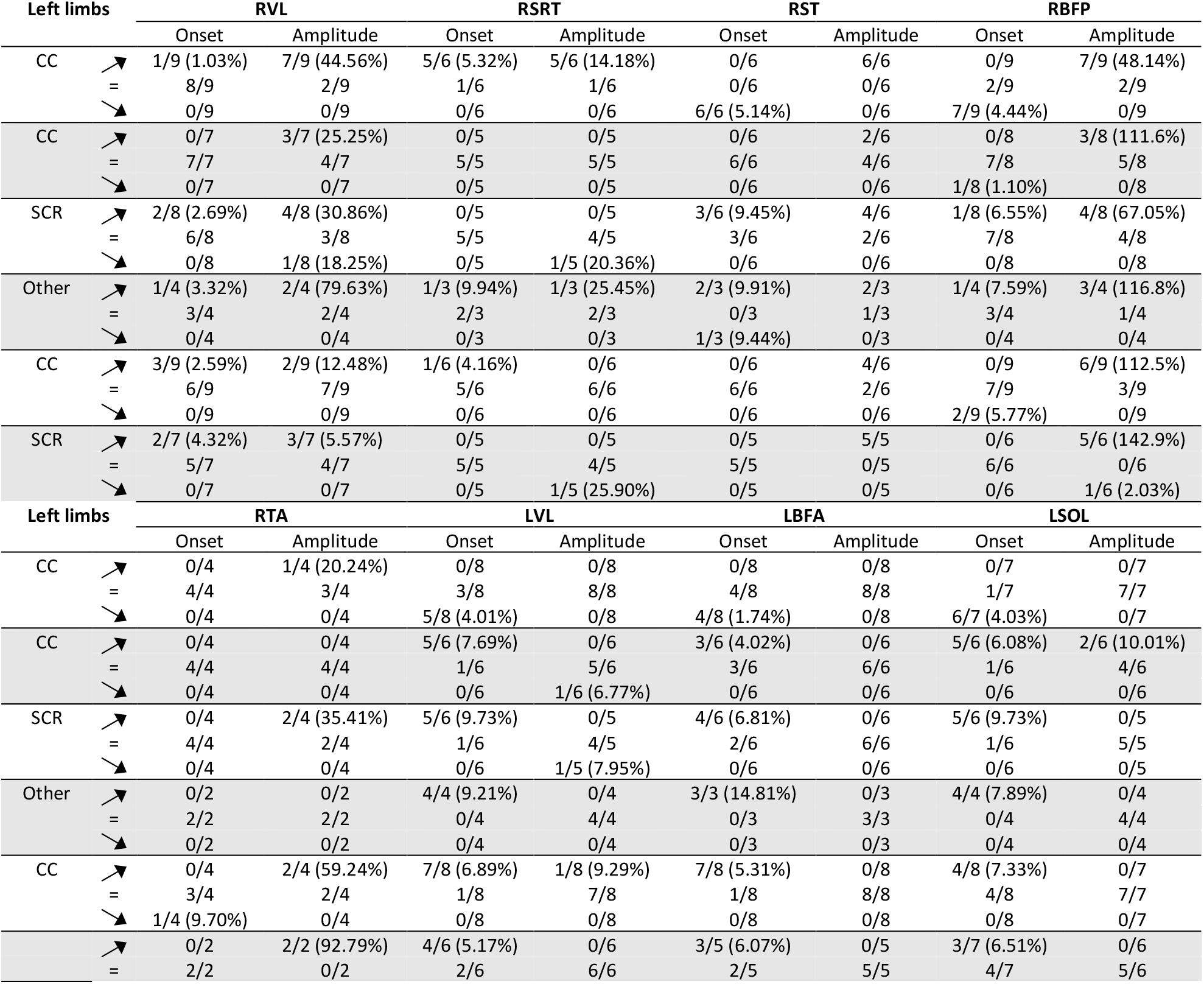
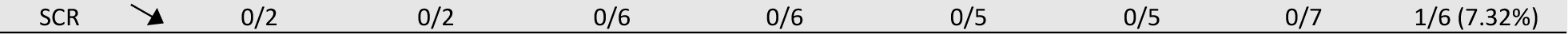
Modulation of EMG burst onset and amplitude before and after lateral hemisection during obstacle negotiation with Left limbs leading. The table shows the number of individual cats/total number of cats with EMGs available for a given muscle. Upward and downward arrows indicate significant increases or decreases, respectively, while the equal sign represents no significant change (two-factor repeated measures ANOVA). The percent value indicates the average of all cats that showed a significant increase or decrease.

**Table 3.** Modulation of EMG burst onset and amplitude before and after lateral hemisection during obstacle negotiation with Right limbs leading. The table shows the number of individual cats/total number of cats with EMGs available for a given muscle. Upward and downward arrows indicate significant increases or decreases, respectively, while the equal sign represents no significant change (two-factor repeated measures ANOVA). The percent value indicates the average of all cats that showed a significant increase or decrease.

| Right limbs |  | RVL |  | RSRT |  | RST |  | RTA |  |
| --- | --- | --- | --- | --- | --- | --- | --- | --- | --- |
|  |  | Onset | Amplitude | Onset | Amplitude | Onset | Amplitude | Onset | Amplitude |
| CC | ↗ | 0/9 | 2/9 (11.05%) | 0/6 | 6/6 (32.53%) | 0/5 | 5/6 (129.1%) | 0/4 | 4/4 (94.05%) |
|  | = | 7/9 | 7/9 | 3/6 | 0/6 | 1/5 | 1/6 | 1/4 | 0/4 |
|  | ↘ | 2/9 (2.52%) | 0/9 | 3/6 (5.42%) | 0/6 | 4/5 (4.44%) | 0/6 | 3/4 (4.73%) | 0/4 |
| CC | ↗ | 0/6 | 0/6 | 0/5 | 1/5 (51.09%) | 0/5 | 0/5 | 0/3 | 0/3 |
|  | = | 6/6 | 6/6 | 5/5 | 4/5 | 5/5 | 5/5 | 3/3 | 3/3 |
|  | ↘ | 0/6 | 0/6 | 0/5 | 0/5 | 0/5 | 0/5 | 0/3 | 0/3 |
| SCR | ↗ | 0/7 | 1/7 (6.19%) | 0/5 | 1/5 (23.17%) | 0/6 | 3/6 (179.5%) | 0/4 | 3/4 (75.63%) |
|  | = | 6/7 | 6/7 | 3/5 | 4/5 | 3/6 | 3/6 | 1/4 | 1/4 |
|  | ↘ | 1/7 (2.43%) | 0/7 | 2/5 (9.22%) | 0/5 | 3/6 (6.11%) | 0/6 | 3/4 (9.15%) | 0/4 |
| Other | ↗ | 0/4 | 0/4 | 0/5 | 0/5 | 0/4 | 0/4 | 0/1 | 0/1 |
|  | = | 4/4 | 4/4 | 5/5 | 5/5 | 4/4 | 4/4 | 1/1 | 1/1 |
|  | ↘ | 0/4 | 0/4 | 0/5 | 0/5 | 0/4 | 0/4 | 0/1 | 0/1 |
| CC | ↗ | 0/9 | 0/8 | 0/5 | 3/5 (45.40%) | 0/5 | 0/5 | 0/3 | 0/3 |
|  | = | 9/9 | 8/8 | 3/5 | 2/5 | 5/5 | 5/5 | 3/3 | 3/3 |
|  | ↘ | 0/9 | 0/8 | 2/5 (7.52%) | 0/5 | 0/5 | 0/5 | 0/3 | 0/3 |
| SCR | ↗ | 0/8 | 0/7 | 0/5 | 1/6 (40.88%) | 0/6 | 6/6 (130.91%) | 0/4 | 4/4 (92.08%) |
|  | = | 8/8 | 7/7 | 1/5 | 5/6 | 6/6 | 0/6 | 1/4 | 0/4 |
|  | ↘ | 0/8 | 0/7 | 4/5 (7.62%) | 0/6 | 0/6 | 0/6 | 3/4 | 0/4 |
| Right limbs |  | LBFA |  | LSOL |  | LSRT |  | LST |  |
|  |  | Onset | Amplitude | Onset | Amplitude | Onset | Amplitude | Onset | Amplitude |
| CC | ↗ | 0/8 | 1/8 (38.74%) | 0/7 | 0/7 | 6/7 (8.98%) | 6/7 (33.17%) | 0/9 | 8/8 (140.7%) |
|  | = | 3/8 | 7/8 | 5/7 | 5/7 | 1/7 | 1/7 | 6/9 | 0/8 |
|  | ↘ | 5/8 (5.46%) | 0/8 | 2/7 (5.03%) | 2/7 (11.37%) | 0/7 | 0/7 | 3/9 (5.25%) | 0/8 |
| CC | ↗ | 0/5 | 0/5 | 0/5 | 0/4 | 0/6 | 0/6 | 0/8 | 0/8 |
|  | = | 5/5 | 5/5 | 5/5 | 4/4 | 6/6 | 6/6 | 8/8 | 8/8 |
|  | ↘ | 0/5 | 0/5 | 0/5 | 0/4 | 0/6 | 0/6 | 0/8 | 0/8 |
| SCR | ↗ | 0/7 | 3/7 (72.23%) | 0/7 | 1/6 (15.15%) | 0/7 | 2/8 (24.69%) | 0/8 | 5/7 (238.8%) |
|  | = | 4/7 | 4/7 | 6/7 | 5/6 | 7/7 | 0/8 | 5/8 | 2/7 |
|  | ↘ | 3/7 (7.77%) | 0/7 | 1/7 (7.72%) | 0/6 | 0/7 | 0/8 | 3/8 (7.09%) | 0/7 |
| Other | ↗ | 0/3 | 0/3 | 0/3 | 0/2 | 0/6 | 0/6 | 0/7 | 3/7 (287.3%) |
|  | = | 3/3 | 3/3 | 3/3 | 2/2 | 6/6 | 6/6 | 6/7 | 4/7 |
|  | ↘ | 0/3 | 0/3 | 0/3 | 0/2 | 0/6 | 0/6 | 1/7 (9.45%) | 0/7 |
| CC | ↗ | 0/7 | 0/7 | 0/8 | 1/7 (18.36%) | 0/6 | 0/6 | 0/8 | 5/8 (78.15%) |
|  | = | 4/7 | 7/7 | 5/8 | 6/7 | 5/6 | 5/6 | 7/8 | 3/8 |
|  | ↘ | 3/7 (4.89%) | 0/7 | 3/8 (6.20%) | 0/7 | 0/6 | 1/6 (22.83%) | 0/8 | 0/8 |
| SCR | ↗ | 0/7 | 3/7 (86.96%) | 0/7 | 0/6 | 1/7 (8.66%) | 4/7 (43.55%) | 1/8 (4.79%) | 6/8 (148.52%) |
|  | = | 4/7 | 4/7 | 5/7 | 6/6 | 6/7 | 3/7 | 6/8 | 2/8 |
|  | ↘ | 3/7 (9.22%) | 0/7 | 2/7 (7.39%) | 0/6 | 0/7 | 0/7 | 1/8 (5.77%) | 0/8 |

For Left limbs leading (Table 2), the muscle strategy to allow the right hindlimb to step over the obstacle for CC in most intact cats consisted of increasing the amplitude of muscles that flex the knee and extend the hip, such as RST (6/6 cats) and RBFP (7/9 cats). RST (6/6 cats) and RBFP (7/9 cats) had also an earlier onset. We also observed a later onset (5/6 cats) and increased amplitude (5/6 cats) of the hip flexor RSRT. The amplitude of the right VL, a knee extensor, also increased in the obstacle step (7/9 cats), possibly to provide greater propulsion to step over the obstacle. In the contralateral hindlimb, we observed an earlier onset of LSOL (6/7 cats). After hemisection, muscle activation strategies were strikingly different. For CC at weeks 1-2, the amplitude of RST (2/6 cats), RBFP (3/8 cats) and RSRT (0/5 cats) did not increase in the obstacle step for all cats. Fewer cats also showed an increase in RVL (3/7 cats). Instead, we saw a delayed onset for LVL (5/6 cats), LBFA (3/6 cats) and LSOL (5/6 cats). At weeks 7-8 for CC, we observed a return of the increase in amplitude for RST (4/6 cats) and RBFP (6/9 cats) but not so much in RVL (2/9 cats). Compared to weeks 1-2, we observed more cats displaying a delayed onset of contralateral extensors, including LVL (7/8 cats), LBFA (7/8 cats) and LSOL (4/8 cats). The SCR at weeks 1-2 was mainly characterized by increased amplitude of RST (4/6 cats), RBFP (4/8 cats) and RTA (2/4 cats) as well as RVL (4/8 cats). We also observed a delayed onset of contralateral extensors, such as LVL (5/6 cats), LBFA (4/6 cats) and LSOL (5/6 cats). The SCR at weeks 7-8 was similar, with an increase in the amplitude of RST (5/5 cats), RBFP (5/6 cats), RTA (2/2 cats) and RVL (3/7 cats). The delayed onset for contralateral extensors was also maintained in about half the animals for LVL (4/6 cats), LBFA (3/5 cats), and LSOL (3/7 cats). For the Other type of negotiation at weeks 1-2, we only observed a significant increase in the amplitude of RBFP (3/4 cats) and a delayed onset of LVL (4/4 cats) and LBFA (3/3 cats).

For Right limbs leading (Table 3), activation strategies for flexor muscles were similar as Left limbs leading for CC in the intact state with an increase in the amplitude of RST (5/6 cats), RSRT (6/6 cats) and RTA (4/4 cats). RSRT (3/6 cats) and RST (4/5 cats) and RTA (3/4 cats) also an earlier onset in the intact state. Using the RVL for propulsion by increasing its amplitude before stepping over the obstacle was less frequent (2/9 cats). At weeks 1-2 and 7-8 after hemisection, the increase in amplitude for flexors was absent for CC, except for RSRT (1/5 and 3/5 cats at weeks 1-2 and 7-8, respectively). The earlier onset of RSRT, RST and RTA was also absent. The SCR at weeks 1-2 was mainly characterized by increased amplitude of RST (3/6 cats) and RTA (3/4 cats) and earlier onsets for RSRT (2/5 cats), RST (3/6 cats) and RTA (3/4 cats). We also observed an increase in the amplitude of contralateral extensors, such as LBFA (3/7 cats) and LSOL (1/6 cats). At weeks 7-8, the increase in the amplitude of RST (6/6 cats) and RTA (4/4) for SCR became more consistent while the increase in LBFA remained similar (3/7 cats). We observed earlier onsets for RSRT (4/5 cats) and RTA (3/4 cats) but not for RST. For the Other type of negotiation at weeks 1-2, we observed no significant modulation of onsets or amplitudes for selected muscles.

In Right limbs leading, we also have information when the left hindlimb stepped over the obstacle for LST and LSRT. For CC in the intact state, we observed an increase in the amplitude of LST (8/8 cats) and LSRT (6/7 cats). We also observed an earlier onset for LST (3/9 cats) and a delay for LSRT (6/7). For CC at weeks 1-2 after hemisection, the increase in the amplitude of LST and LSRT and the delayed onset in LSRT were not present in all cats. The SCR at weeks 1-2 was mainly characterized by increased amplitude of LST (5/7 cats). For Other, LST increased in amplitude in some cats (3/7 cats) with only one cat showing an earlier onset (1/7 cats). At 7-8 weeks after the hemisection, CC was characterized by an increased amplitude for LST (5/8 cats) only, with an earlier onset observed for LSRT in only one cat (1/6 cats). For SCR at weeks 7-8, we found an increase in amplitude for LST (6/8 cats) and LSRT (4/7 cats) in most cats. Changes in onset were rare.

## Discussion

In the present study, we showed that intact cats easily stepped over an obstacle without contact, or complete clearance. However, after a mid-thoracic lateral hemisection, cats displayed different types of obstacle negotiations of their ipsilesional hindlimb, using different neuromechanical strategies, even with complete clearance. Below, we explain these findings in terms of neuroplastic changes within the central nervous system and by using an optimal control framework.

### The appearance of different negotiation types reflects a loss in anticipatory control and a decrease in spinal neuronal excitability

Intact cats clear an obstacle by mainly flexing the knee followed to a lesser degree by flexing the ankle with Left and Right limbs leading (Fig. 5). This is consistent with previous results in cats [29] and humans [28]. After hemisection, the knee flexion strategy was maintained in both conditions whereas an additional ankle flexion occurred with Left limbs leading. A SCR appeared when the cat’s hindpaw contacted the obstacle, due to an inability to sufficiently activate muscles that flex the knee before the obstacle and/or because of improper timing of joint flexions (Fig. 6). Earlier flexion of the hip after hemisection may bring the limb forward prematurely before sufficient elevation occurs, producing contact with the obstacle. With Right limbs leading, an earlier knee flexion may compensate for earlier hip flexion to allow sufficient elevation in some trials.

Studies in cats and humans have shown that the SCR is triggered mainly by cutaneous afferents, particularly from the superficial peroneal nerve that innervates the foot dorsum [10–14,16,17,32,43–47]. In our study, all cats showed a SCR in the first two weeks after hemisection. Doperalski et al. (2011) also found that a SCR appeared between 2 and 4 weeks after hemisection. Studies have reported that the SCR and spinal reflexes in general are depressed in the ipsilesional hindlimb for a few days after hemisection but come back in the first two weeks as locomotion also recovers [48,49]. The SCR probably shares some circuits with the withdrawal reflex, which becomes more variable after a lateral hemisection, with greater variability in latency [50].

The absence of SCR following contact, or the Other negotiation type, is surprising because the SCR can be triggered in spinal cats [19,20]. Why does the SCR fail to appear following some obstacle contacts? The lack of an SCR could be due to a decrease in spinal neuronal excitability, which can persist for several weeks after hemisection. Studies have also shown that supraspinal pathways, such as the rubrospinal tract, facilitate the SCR in cats [51–53]. The lack of SCR could also be due to the area of the hindpaw that makes contact with the obstacle, for example the toes versus the foot dorsum [54]. The decrease in Other from weeks 1-2 to weeks 7-8 after hemisection coupled to an increase in complete clearance reflects a partial recovery of anticipatory control.

### Neuromechanical strategies when negotiating obstacles before after hemisection

Cats have a preferential leading limb when grabbing food, going down stairs or when stepping into a litter box [55] but not when it comes to negotiating an obstacle, as shown here and in a previous study [25]. Cats also cannot be categorized as mainly right- or left-handed. The lack of side preference persisted at weeks 1-2 after hemisection but at weeks 7-8, we observed a clear preference for Left limbs leading, the contralesional side (> 60% of trials). Functionally, this is a more stable position in terms of balance because the contralesional limb has already cleared the obstacle and is in a stable support position to allow the ipsilesional limb to step over the obstacle. This might be a conscious decision/strategy made by cats.

To further test this hypothesis, we determined the redistribution of weight support after hemisection when negotiating obstacles by measuring support periods (Fig. 7). Generally, the triple support involving both hindlimbs and the left forelimb decreased after hemisection as did both diagonal supports for most types of negotiations. The absence of a decrease in the diagonal support involving the right hindlimb and left forelimb did not decrease for Other at weeks 1-2 after hemisection. Diagonal support periods are the least stable support when stepping in the forward direction [56] and this lack of a decrease after hemisection could have contributed to the appearance of Other. In contrast, homolateral support periods increased after hemisection, particularly on the contralesional side, where it approximately doubled in proportion for all negotiation types. Thus, cats shift a considerable portion of their weight to the contralesional side after hemisection to negotiate obstacles, consistent with an effort to maximize dynamic balance.

### Stepping over an obstacle as an optimization problem

Clearing an obstacle relies on anticipatory control, or planning [3,57–62]. Intact cats increased their speed when stepping over the obstacle in both Left and Right limbs leading conditions (Fig. 8). Humans also increase their speed before stepping on a curb based on optimization criteria to conserve momentum and minimize energy expenditure by modulating push-off work [63]. Most cats increased the amplitude of RVL, a knee extensor, in the stance phase that preceded stepping over the obstacle in the Left limbs leading condition, consistent with an increase in push-off work. This increase in speed and in RVL amplitude was lost after hemisection.

We also showed that ipsilesional hindlimb flexion was exaggerated after clearing the obstacle with Left limbs leading at both time points after hemisection (Fig. 3B). There is no functional benefit to the exaggerated flexion observed after complete clearance because it increases energy expenditure [64] and potentially destabilizes the body. We can explain the exaggerated flexion following clearance in the Left limbs leading condition using the framework of optimal feedback control theory [65,66]. In optimal feedback control, a state estimator receives sensory information and sends signals to a controller, which controls a policy or a cost function, in our case energy expenditure. The controller sends an efferent copy of the motor commands to the state estimator and to a forward internal model that predicts the state, allowing the controller to perform online corrections based on all available information. After hemisection, we propose that proprioceptive information cannot reach the state estimator and/or the forward internal model, located in the cerebellum for example [65], and the controller located in the spinal cord [67] cannot properly correct ipsilesional hindlimb trajectory and minimize energy expenditure. Future studies can apply computational approaches to test this hypothesis.

### Neuroplastic changes and functional recovery

In an intact system, descending pathways from the brain and propriospinal pathways from the cervical cord project to lumbar interneurons controlling locomotion, such as those of the spinal locomotor CPG, while proprioceptive and tactile afferent inputs from the limbs terminate in spinal circuits and ascend to the brain (Fig. 10, left panel). In intact cats, studies have shown contributions from rubro- and corticospinal tracts in negotiating obstacles [6–9,24,68]. A lateral thoracic spinal hemisection abolishes all descending and ascending pathways on one side of the cord (Fig. 10, mid panel). We can attribute the smaller proportion of complete clearance at 1-2 weeks after hemisection to the loss of rubro- and corticospinal pathways, a decrease in spinal neuronal excitability and/or to a loss of ascending proprioceptive information. Spared connections from the contralesional corticospinal tract, which has extensive commissural projections throughout the spinal cord in cats, can explain the ability to maintain obstacle clearance after hemisection [69,70].

**Figure 10.**
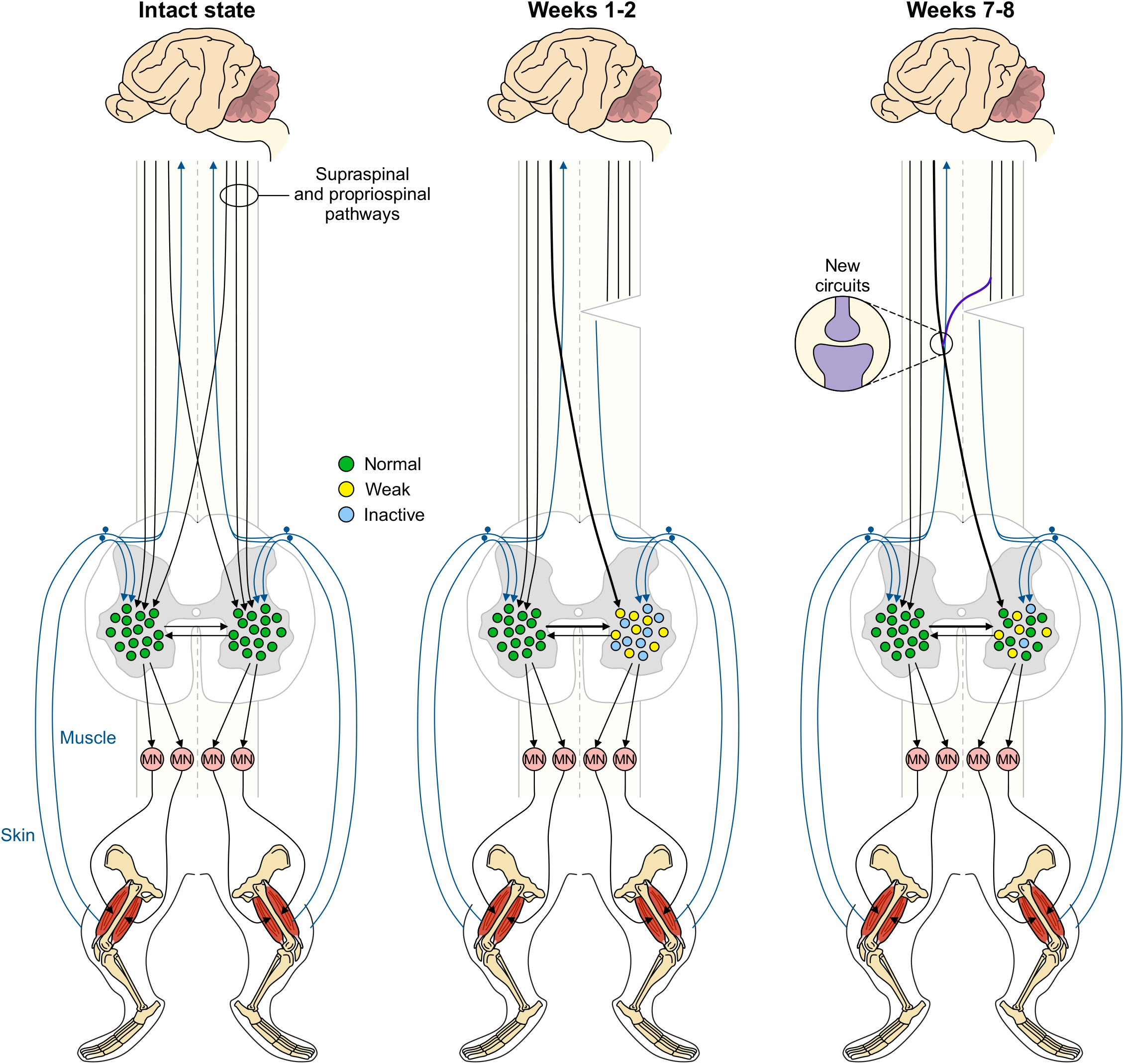
Neuroplastic changes after hemisection and potential mechanisms of recovery. In intact the state, descending supraspinal and propriospinal pathways reach spinal interneurons of the lumbar cord. These spinal interneurons control spinal motoneurons controlling limb muscles. Pathways transmitting signals from proprioceptive and cutaneous afferents ascend to the brain and project locally to spinal interneurons and motoneurons (not shown). At weeks 1-2 after hemisection, descending and ascending spinal pathways are disrupted and some spinal interneurons on the right side display weak excitability or a lack of activity. Strengthened descending pathways from the contralesional side and commissural pathways from the spinal interneuronal network controlling the left hindlimb are strengthened, as shown by thicker lines. At weeks 7-8 after hemisection, spinal interneurons on the right side recover their excitability and new descending propriospinal pathways can form.

The recovery of complete clearance from early to late time points after hemisection can be due to several factors, such as a return in spinal neuronal excitability and new or strengthened pathways from the brain, for instance the contralesional corticospinal tract (Fig. 10, right panel). Projections from corticospinal axons increase in the spinal cord caudal to a lesion in rats [71,72]. While new or strengthened contralesional pathways are not necessary for the recovery of locomotion after incomplete SCI [49,73], they might be essential for the recovery of functions requiring voluntary/anticipatory control, such as obstacle negotiation. Other descending pathways may also contribute, such as the reticulospinal [74,75] and rubrospinal [76] tracts.

Descending pathways from the brain also synapse on long and short propriospinal neurons in the cervical cord that then project to the lumbar region [77]. After incomplete SCI, such as a lateral hemisection, studies have shown that new or strengthened propriospinal pathways can transmit commands from the brain to the lumbar cord [78–85]. Although reorganized propriospinal circuits might bypass the lesion site by projecting to the other side [86,87], whether these new connections make functional and meaningful contributions to locomotor control is less clear.

Neuroplastic changes caudal to the lesion likely play an important role in the recovery of obstacle clearance. Studies in cats have shown that the spinal locomotor network becomes more autonomous following an incomplete SCI and that sensorimotor interactions are altered [88–91]. Spinal interneurons can form new connections [92] and excitability returns, with some neurons showing constitutive activity [81] These changes within neuronal circuits of the lumbosacral cord make them more responsive to descending inputs, thus facilitating voluntary commands. Contralesional spinal circuits might also facilitate the recovery of anticipatory control through strengthened commissural interactions [93].

## Conclusions

The ability to perform various locomotor tasks, particularly those involving voluntary or anticipatory control remains a challenge for people with SCI. In the present study, we showed partial maintenance of anticipatory control after a mid-thoracic lateral hemisection in cats and some recovery over time, albeit incomplete. Future studies will need to address how we can promote neuroplastic changes to restore complete clearance of obstacles after incomplete SCI.

## Materials and methods

### Animals and ethical information

The Animal Care Committee of the Université de Sherbrooke approved all procedures in accordance with the policies and directives of the Canadian Council on Animal Care (Protocol 442-18). In the present study, we used ten adult cats (>1 year of age at the time of experimentation), five females and five males, with a mass between 3.8 kg and 6.1 kg. We followed the ARRIVE guidelines for animal studies [30]. To reduce the number of animals used in research, cats participated in other studies to answer different scientific questions [31,32].

### Surgical procedures and electrode implantation

We performed surgeries under aseptic conditions with sterilized instruments in an operating room. Prior to surgery, the cat was sedated with an intramuscular injection of butorphanol (0.4 mg/kg), acepromazine (0.1 mg/kg) and glycopyrrolate (0.01 mg/kg) and inducted with another intramuscular injection of ketamine and diazepam (0.05 ml/kg) in a 1:1 ratio. The fur overlying the back, stomach, forelimbs and hindlimbs was shaved and the skin was cleaned with chlorhexidine soap. The cat was then anesthetized with isoflurane (1.5-3%) and O_2_ using a mask for a minimum of 5 minutes and then intubated with a flexible endotracheal tube. Isoflurane concentration was confirmed and adjusted throughout the surgery by monitoring cardiac and respiratory rates, by applying pressure to the paw to detect limb withdrawal and by assessing muscle tone. A rectal thermometer was used to monitor body temperature and keep it within physiological range (37 ± 0.5°C) using a water-filled heating pad placed under the animal and an infrared lamp positioned ∼ 50 cm above the cat. Cats received a continuous infusion of lactated Ringer’s solution (3 ml/kg/h).

We directed pairs of Teflon-insulated multistrain fine wires (AS633; Cooner Wire, Chatsworth, CA, USA) subcutaneously from two head-mounted 34-pin connectors (Omnetics Connector Corporation, Minneapolis, MN, USA) and sewn them into the belly of selected forelimb and hindlimb muscles for bipolar recordings, with 1-2 mm of insulation stripped from each wire. The head connector was secured to the skull using dental acrylic and six screws (Fig. 11A). We verified electrode placement by electrically stimulating each muscle through the appropriate head connector channel.

**Figure 11.**
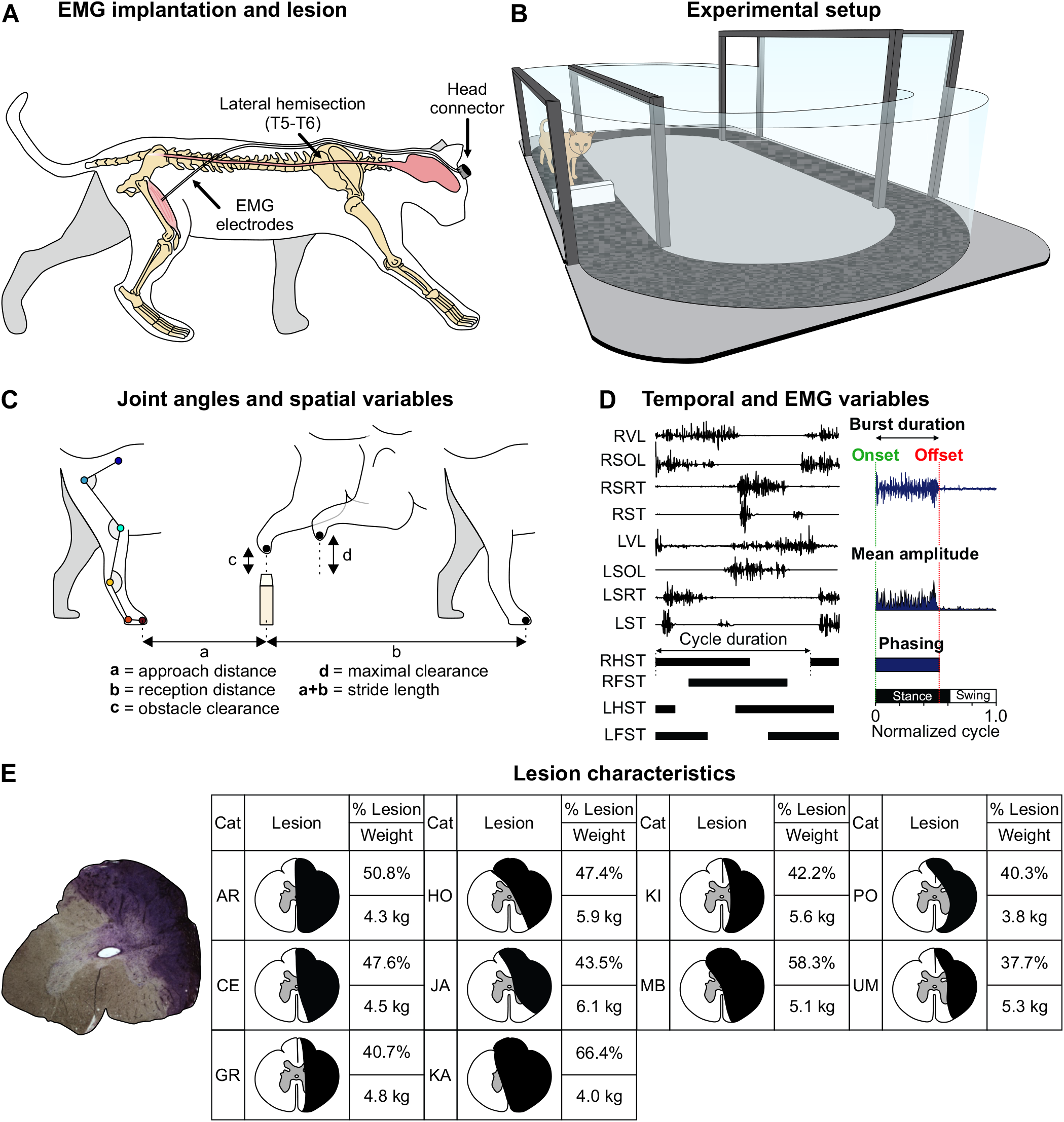
Experimental design, data analysis and lesion characteristics. **A**: Experimental design showing EMG electrodes directed subcutaneously to muscles from a head connector and the site of spinal lesion. **B**: Walkway with an obstacle in the straight path. **C**: Joint angles and spatial variables. **D**: Measures of EMG variables. EMG activity is shown for 4 muscles of the left (L) hindlimb and 4 muscles of the right (R) hindlimb, VL, vastus lateralis; SOL, soleus; SRT, anterior sartorius; ST, semitendinosus. Thick horizontal lines represent right hindlimb (RHST), right forelimb (RFST), left hindlimb (LHST) and left forelimb (LFST) stances phases. **E**: Spinal lesion site with Cresyl violet staining (Cat JA). The darker area is the lesioned area. Table shows lesion extent for all cats.

At the end of surgery, we injected an antibiotic (Cefovecin, 0.1 ml/kg) subcutaneously and taped a transdermal fentanyl patch (25 µg/h) to the back of the animal 2-3 cm rostral to the base of the tail. We injected buprenorphine (0.01 mg/kg), a fast-acting analgesic, subcutaneously during surgery and ∼ 7 h after. Following surgery, we placed the cats in an incubator until they regained consciousness. After the experiments, under general anesthesia, cats received a lethal dose of pentobarbital (120 mg/kg) through the left or right cephalic vein and we collected spinal cord tissue for histological analysis.

### Lateral hemisection

A small laminectomy was performed between the fifth and sixth thoracic vertebrae. After exposing the spinal cord, lidocaine (Xylocaine, 2%) was applied topically and injected intraspinally on the right side of the cord, which was then hemisected laterally from the midline with surgical scissors. Hemostatic material (Spongostan) was inserted within the gap, and muscles and skin were sewn back to close the opening in anatomic layers. The same pre- and post-operative treatments were administered as for the implantation surgery. In the days following the hemisection, cats were carefully monitored for voluntary bodily functions. The bladder was expressed manually if needed.

### Experimental protocol

We trained cats to step on an overground walkway with a straight path 207 cm long and 32 cm wide between two Plexiglas with food and affection as reward (Fig. 11B). The walkway is oval-shaped and the animals have sufficient room to turn around at the two ends of the straight path. The surface of the walkway is made of black rubberized material. To assess obstacle negotiation, cats stepped in the walkway with an obstacle placed in the middle of the straight path. The three obstacles used were white plastic objects 2.5 cm wide and 1, 5 and 9 cm in height that could easily be knocked over if contacted by the cat’s limb to avoid injury. From the first training session, cats negotiated all obstacle heights without contact. Experiments began after a minimum of 2 weeks of familiarization where cats performed sessions lasting 20-30 minutes, 3 times a week.

Each cat performed one session that consisted of negotiating the three obstacle heights. Cats were given a few seconds of rest and a food reward between each negotiation (a trial) to obtain ∼ 20 trials for each obstacle height where we could analyze the step before, during and after crossing the obstacle. A session lasted ∼ 25-30 minutes. Before and after hemisection, cats stepped at a self-selected speed. The trials where cats were running, jumping or pausing between different steps were not analyzed.

### Data acquisition and analysis

We collected kinematic and EMG data before, 1-2 weeks and 7-8 weeks after hemisection. Two cameras (Basler AcA640-100g) captured videos of the left and right sides at 60 frames per second with a spatial resolution of 640×480 pixels. A custom-made program (LabView) acquired the images and synchronized acquisition with EMG data. We analyzed each video offline using a deep-learning approach, DeepLabCut™ [33,34], as recently described [31,32,35].

We determined the contact and liftoff of each limb by visual inspection. We defined contact as the first frame where the paw made visible contact with the walking surface while liftoff corresponded to the frame with the most caudal displacement of the toe. Based on contacts and liftoffs for each limb, we measured individual periods of support (double, triple and quad) and expressed them as a percentage of cycle duration [31,36]. During a normalized cycle, defined from successive right hindlimb contacts, we identified nine periods of limb support [37–40]. We measured various spatial parameters (Fig. 11C), including right hindlimb stride length, defined as the distance traveled by the limb between two consecutive contacts. We measured the animal’s speed during a cycle by dividing the horizontal displacement of the right hip between two consecutive right hindlimb contacts by cycle duration. We obtained the height of the paw as it travelled over the obstacle by measuring the distance between the base of the fifth metatarsal and the top of the obstacle when the paw is directly above the obstacle (obstacle clearance) and when it reaches its maximum height (maximal clearance). We measured two other distances horizontally from the base of the fifth metatarsal and the obstacle. The approach distance was measured at the time of right hindlimb contact before the obstacle. The reception distance was measured at right hindlimb contact after the obstacle. We measured right hip, knee and ankle angles throughout the step cycle.

EMG signals were pre-amplified (x10, custom-made system), bandpass filtered (30-1000Hz) and amplified (100-5000x) using a 16-channel amplifier (AM Systems Model 3500, Sequim, WA, USA). As we implanted more than 16 muscles per cat, we obtained data for each obstacle height twice, one for each connector, as our data acquisition system does not currently allow us to record more than 16 channels simultaneously. EMG data were digitized (2000 Hz) with a National Instruments card (NI 6032E), acquired with a custom-made acquisition software and stored on a computer. Although several muscles were implanted in the fore- and hindlimbs, we focused our analysis on ten muscles of the left (L) and right (R) hindlimbs: vastus lateralis (knee extensor, LVL, n = 8; RVL, n = 9), biceps femoris anterior (hip extensor, LBFA, n = 8; RBFA, n = 9), lateral gastrocnemius (ankle extensor/knee flexor, LLG, n = 7; RLG, n = 8), medial gastrocnemius (ankle extensor/knee flexor, LMG, n = 7; RMG, n = 7), soleus (ankle extensor, LSOL n = 8; RSOL, n = 9), anterior sartorius (hip flexor/knee extensor, LSRT, n = 7; RSRT, n = 6), the iliopsoas (hip flexor, LIP, n = 3; RIP, n = 3), the biceps femoris posterior (hip extensor/knee flexor, LBFP, n = 8; RBFP, n = 8), the semitendinosus (hip extensor/knee flexor, LST, n = 9; RST, n = 6) and the tibialis anterior (ankle flexor, RTA, n = 4). Burst onsets and offsets were determined visually by the same experimenter (Lecomte) from the raw EMG waveforms using a custom-made program. Burst duration was determined from onset to offset while mean EMG amplitude was measured by integrating the full-wave rectified EMG burst from onset to offset and dividing it by its burst duration (Fig 1D). Joint angles were low-pass filtered (fourth-order Butterworth filter, zero-lag, cutoff frequency 6Hz).

### Histology

To confirm the extent of the spinal lesion, we performed histology. Following euthanasia, we dissected a 2-cm long segment of the spinal cord centered around the injury site and placed it in 25 mL of 4% paraformaldehyde solution (PFA in 0.1M PB, 4°C). After five days, the spinal cord was cryoprotected in PB 0.2M containing 30% sucrose for 72 h at 4°C. We then sliced the spinal cord into 50 µm coronal sections using a cryostat (Leica CM1860, Leica Biosystems Inc, Concord, ON, Canada). Sections were mounted on gelatinized slides, dried overnight, and stained with 1% Cresyl violet for 12 minutes. The slides were then washed once in water for 3 minutes before being dehydrated in successive baths of ethanol 50%, 70% and 100%, 5 minutes each. After a final 5 minutes bath in xylene, slides were mounted with dibutylphthalate polystyrene xylene (DPX, Sigma-Aldrich Canada) and dried before being scanned using a Nanozoomer (Hamamastu Corporation, Bridgewater Township, NJ, USA). We then performed qualitative and quantitative evaluations of the lesioned area (**Fig. 11E**).

### Statistical analysis

We performed statistical tests with IBM SPSS Statistics 20.0 (IBM Corp., Armonk, NY, US). We separated trials with left and right limbs leading. When the right forelimb stepped over the obstacle first, it was followed by the left forelimb, right hindlimb and left hindlimb. We termed these trials Right limb leading. The same applies to Left limb leading (left forelimb followed by right forelimb, left hindlimb and right hindlimb). For each animal and a given obstacle height, we averaged Left and Right limb leading trials separately. We performed a two-factor mixed-effects model ANOVA with the cat as the experimental unit to determine main effects and possible interactions on dependent variables for the following factors: 1) time point x obstacle height, 2) side x obstacle height, 3) negotiation x obstacle height, 4) step x obstacle height and 5) clearance x obstacle height. For each two-factor ANOVA, the interaction between the two parameters determines if one factor influences the other. For EMG data, we performed a two-factor repeated measures ANOVA for individual cats to determine individualized strategies. The normality of each variable was assessed by the Shapiro-Wilk test. Although we did not correct for multiple comparisons to avoid Type II errors [see 41,42 for a discussion], we used a confidence interval of 99% or a P ≤ 0.01 to determine significance. This increases the probability that significant differences correspond to robust and physiological/functional changes in the gait pattern. All P values for main effects of statistical tests can be found in S3-S8 Tables.

## Acknowledgments

We thank Philippe Drapeau for providing data acquisition and analysis software, developed in the Rossignol and Drew laboratories.

## Supporting information

**S1 Table. Modulation of EMG phasing and amplitude before and after lateral hemisection during obstacle negotiation with Left limbs leading**. The table shows the number of individual cats/total number of cats with EMGs available for a given muscle. Upward and downward arrows indicate significant increases or decreases, respectively, while the equal sign represents no significant change (two-factor repeated measures ANOVA). The percent value indicates the average of all cats that showed a significant increase or decrease.

**S2 Table. Modulation of EMG phasing and amplitude before and after lateral hemisection during obstacle negotiation with Right limbs leading**. The table shows the number of individual cats/total number of cats with EMGs available for a given muscle. Upward and downward arrows indicate significant increases or decreases, respectively, while the equal sign represents no significant change (two-factor repeated measures ANOVA). The percent value indicates the average of all cats that showed a significant increase or decrease.

**S3 Table. P values for each negotiation type and side preference**. P values comparing time and heights are indicated (mixed-effects model ANOVA) for each negotiation type (Fig. 1B). P values comparing side and heights are indicated (mixed-effects model ANOVA) for each type of negotiation (Fig. 1C).

**S4 Table. P values for distances from the obstacles and difference of clearance**. P values comparing each negotiation type and heights after hemisection to complete clearance in intact cats are indicated (mixed-effects model ANOVA) for approach distance, reception distance and difference of clearance (Fig. 3).

**S5 Table. P values for angle excursions**. P values comparing steps and heights are indicated (mixed-effects model ANOVA) for each type of negotiation for hip, knee and ankle angular excursions (Fig. 5).

**S6 Table. P values for timing of joint flexions**. P values comparing each negotiation type and obstacle heights to complete clearance in intact cats are indicated (mixed-effects model ANOVA) for the timing of hip, knee and ankle joint flexion (Fig. 6).

**S7 Table. P values for support periods**. P values comparing each negotiation type and obstacle heights to complete clearance in intact cats are indicated (mixed-effects model ANOVA) for support periods (Fig. 7).

**S8 Table. P values for speed and stride length**. P values comparing the control and obstacle steps and heights are indicated (mixed-effects model ANOVA) for each type of negotiation for speed and stride length (Fig. 8).

