## Supplementary tables for "Neuromechanical strategies for obstacle negotiation during overground locomotion following an incomplete spinal cord injury in adult cats"

| Left limbs leading | | RBFA | | RMG | | RLG | | RSOL | |
|---|---|---|---|---|---|---|---|---|---|
| | | Onset | Amplitude | Onset | Amplitude | Onset | Amplitude | Onset | Amplitude |
| CC INTACT | ↗ | 0/9 | 3/9 (65.84%) | 0/7 | 0/7 | 0/8 | 0/8 | 2/9 (2.78%) | 1/9 (8.94%) |
| | = | 9/9 | 6/9 | 7/7 | 7/7 | 8/8 | 8/8 | 7/9 | 8/9 |
| | ↘ | 0/9 | 0/9 | 0/7 | 0/7 | 0/8 | 0/8 | 0/9 | 0/9 |
| CC H1-2 | ↗ | 0/8 | 2/8 (64.36%) | 0/6 | 0/6 | 0/7 | 0/7 | 0/7 | 0/7 |
| | = | 8/8 | 6/8 | 6/6 | 6/6 | 7/7 | 6/7 | 7/7 | 7/7 |
| | ↘ | 0/8 | 0/8 | 0/6 | 0/6 | 0/7 | 1/7 (23.94%) | 0/7 | 0/7 |
| SCR H1-2 | ↗ | 0/8 | 0/8 | 1/6 (1.82%) | 0/6 | 1/6 (6.36%) | 0/6 | 0/8 | 0/7 |
| | = | 8/8 | 8/8 | 5/6 | 6/6 | 5/6 | 6/6 | 8/8 | 6/7 |
| | ↘ | 0/8 | 0/8 | 0/6 | 0/6 | 0/6 | 0/6 | 0/8 | 1/7 (17.04%) |
| Other H1-2 | ↗ | 0/4 | 0/4 | 0/4 | 0/4 | 1/4 (3.08%) | 0/4 | 1/4 (4.33%) | 0/4 |
| | = | 4/4 | 4/4 | 4/4 | 4/4 | 3/4 | 4/4 | 3/4 | 4/4 |
| | ↘ | 0/4 | 0/4 | 0/4 | 0/4 | 0/4 | 0/4 | 0/4 | 0/4 |
| CC H7-8 | ↗ | 2/9 (4.22%) | 4/9 (38.60%) | 0/7 | 0/7 | 0/8 | 0/8 | 1/9 (2.24%) | 1/9 (12.09%) |
| | = | 7/9 | 5/9 | 7/7 | 7/7 | 8/8 | 8/8 | 8/9 | 7/9 |
| | ↘ | 0/9 | 0/9 | 0/7 | 0/7 | 0/8 | 0/8 | 0/9 | 1/9 (6.98%) |
| SCR H7-8 | ↗ | 1/6 (5.03%) | 0/6 | 1/4 (4.72%) | 0/4 | 1/7 (5.87%) | 0/7 | 2/7 (2.08%) | 0/7 |
| | = | 5/6 | 5/6 | 3/4 | 2/4 | 6/7 | 7/7 | 5/7 | 7/7 |
| | ↘ | 0/6 | 1/6 (17.08%) | 0/4 | 2/4 (19.15%) | 0/7 | 0/7 | 0/7 | 0/7 |

| Left limbs leading | | RIP | | LMG | | LLG | |
|---|---|---|---|---|---|---|---|
| | | Onset | Amplitude | Onset | Amplitude | Onset | Amplitude |
| CC INTACT | ↗ | 2/2 (2.39%) | 0/2 | 0/7 | 0/7 | 0/7 | 0/7 |
| | = | 0/2 | 2/2 | 1/7 | 7/7 | 3/7 | 7/7 |
| | ↘ | 0/2 | 0/2 | 6/7 (3.79%) | 0/7 | 4/7 (4.87%) | 0/7 |
| CC H1-2 | ↗ | 0/2 | 0/2 | 1/6 (3.59%) | 0/5 | 2/7 (5.02%) | 0/6 |
| | = | 2/2 | 2/2 | 5/6 | 4/5 | 5/7 | 5/6 |
| | ↘ | 0/2 | 0/2 | 0/6 | 1/5 (9.75%) | 0/7 | 1/6 (9.22%) |
| SCR H1-2 | ↗ | 0/2 | 1/2 (23.39%) | 3/5 (4.30%) | 0/5 | 4/6 (8.61%) | 0/5 |
| | = | 1/2 | 1/2 | 2/5 | 5/5 | 2/6 | 4/5 |
| | ↘ | 1/2 (6.95%) | 0/2 | 0/5 | 0/5 | 0/6 | 1/5 (11.03%) |
| Other H1-2 | ↗ | 0/2 | 0/2 | 4/4 (6.07%) | 0/4 | 3/3 (6.43%) | 0/3 |
| | = | 2/2 | 2/2 | 0/4 | 4/4 | 0/3 | 3/3 |
| | ↘ | 0/2 | 0/2 | 0/4 | 0/4 | 0/3 | 0/3 |
| CC H7-8 | ↗ | 0/2 | 0/2 | 4/7 (3.51%) | 0/7 | 5/7 (4.47%) | 0/7 |
| | = | 2/2 | 2/2 | 3/7 | 6/7 | 2/7 | 7/7 |
| | ↘ | 0/2 | 0/2 | 0/7 | 1/7 (13.98%) | 0/7 | 0/7 |
| SCR H7-8 | ↗ | 0/0 | 0/0 | 1/4 (7.22%) | 0/4 | 4/6 (8.11%) | 1/6 (7.76%) |
| | = | 0/0 | 0/0 | 3/4 | 4/4 | 2/6 | 4/6 |
| | ↘ | 0/0 | 0/0 | 0/4 | 0/4 | 0/6 | 1/6 (12.98%) |

**Table S1. Modulation of EMG burst onset and amplitude before and after lateral hemisection during obstacle negotiation with Left limbs leading.** The table shows the number of individual cats/total number of cats with EMGs available for a given muscle. Upward and downward arrows indicate significant increases or decreases, respectively, while the equal sign represents no significant change (two-factor repeated measures ANOVA). The percent value indicates the average of all cats that showed a significant increase or decrease.

| Right limbs leading | | RBFA | | RMG | | RLG | | RSOL | |
|---|---|---|---|---|---|---|---|---|---|
| | | Onset | Amplitude | Onset | Amplitude | Onset | Amplitude | Onset | Amplitude |
| CC INTACT | ↗ | 0/9 | 4/9 (37.29%) | 0/7 | 3/7 (29.09%) | 0/8 | 4/8 (13.05%) | 1/9 (2.40%) | 0/9 |
| | = | 7/9 | 5/9 | 7/7 | 4/7 | 6/8 | 4/8 | 8/9 | 8/9 |
| | ↘ | 2/9 (2.49%) | 0/9 | 0/7 | 0/7 | 2/8 (0.96%) | 0/8 | 0/9 | 1/9 (13.15%) |
| CC H1-2 | ↗ | 0/6 | 0/6 | 0/4 | 0/4 | 0/8 | 0/8 | 0/6 | 0/6 |
| | = | 6/6 | 6/6 | 4/4 | 4/4 | 8/8 | 8/8 | 6/6 | 6/6 |
| | ↘ | 0/6 | 0/6 | 0/4 | 0/4 | 0/8 | 0/8 | 0/6 | 0/6 |
| SCR H1-2 | ↗ | 0/8 | 0/8 | 1/6 (2.79%) | 1/6 (35.15%) | 0/7 | 0/7 | 0/8 | 1/8 (28.27%) |
| | = | 8/8 | 8/8 | 5/6 | 5/6 | 7/7 | 7/7 | 8/8 | 7/8 |
| | ↘ | 0/8 | 0/8 | 0/6 | 0/6 | 0/7 | 0/7 | 0/8 | 0/8 |
| Other H1-2 | ↗ | 0/3 | 0/3 | 0/2 | 0/2 | 0/6 | 0/6 | 0/4 | 1/4 (44.22%) |
| | = | 3/3 | 3/3 | 2/2 | 2/2 | 6/6 | 6/6 | 4/4 | 3/4 |
| | ↘ | 0/3 | 0/3 | 0/2 | 0/2 | 0/6 | 0/6 | 0/4 | 0/4 |
| CC H7-8 | ↗ | 0/8 | 0/8 | 0/6 | 0/6 | 0/8 | 0/8 | 1/9 (2.82%) | 0/9 |
| | = | 8/8 | 8/8 | 6/6 | 6/6 | 8/8 | 8/8 | 8/9 | 9/9 |
| | ↘ | 0/8 | 0/8 | 0/6 | 0/6 | 0/8 | 0/8 | 0/9 | 0/9 |
| SCR H7-8 | ↗ | 1/8 (4.25%) | 0/8 | 0/7 | 0/7 | 3/8 (3.51%) | 0/7 | 0/8 | 0/8 |
| | = | 7/8 | 8/8 | 7/7 | 7/7 | 5/8 | 7/7 | 8/8 | 8/8 |
| | ↘ | 0/8 | 0/8 | 0/7 | 0/7 | 0/8 | 0/7 | 0/8 | 0/8 |

| Right limbs Leading | | RIP | | RBFP | | LVL | | LMG | |
|---|---|---|---|---|---|---|---|---|---|
| | | Onset | Amplitude | Onset | Amplitude | Onset | Amplitude | Onset | Amplitude |
| CC INTACT | ↗ | 0/2 | 2/2 (48.14%) | 0/9 | 3/9 (42.87%) | 0/8 | 6/7 (39.05%) | 0/7 | 0/7 |
| | = | 1/2 | 0/2 | 4/9 | 6/9 | 4/8 | 1/7 | 2/7 | 7/7 |
| | ↘ | 1/2 (3.19%) | 0/2 | 5/9 (5.16%) | 0/9 | 4/8 (6.23%) | 0/7 | 5/7 (5.08%) | 0/7 |
| CC H1-2 | ↗ | 0/1 | 0/1 | 0/6 | 0/6 | 0/5 | 0/5 | 0/4 | 0/4 |
| | = | 1/1 | 1/1 | 6/6 | 6/6 | 5/5 | 5/5 | 4/4 | 4/4 |
| | ↘ | 0/1 | 0/1 | 0/6 | 0/6 | 0/5 | 0/5 | 0/4 | 0/4 |
| SCR H1-2 | ↗ | 0/2 | 2/2 (52.41%) | 0/8 | 4/8 (124.95%) | 0/7 | 3/7 (48.02%) | 0/6 | 0/6 |
| | = | 2/2 | 0/2 | 5/8 | 4/8 | 4/7 | 4/7 | 2/6 | 6/6 |
| | ↘ | 0/2 | 0/2 | 3/8 (10.05%) | 0/8 | 3/7 (10.03%) | 0/7 | 4/6 (8.78%) | 0/6 |
| Other H1-2 | ↗ | 0/1 | 0/1 | 0/3 | 0/3 | 0/3 | 0/3 | 0/1 | 0/2 |
| | = | 1/1 | 1/1 | 3/3 | 3/3 | 3/3 | 3/3 | 1/1 | 2/2 |
| | ↘ | 0/1 | 0/1 | 0/3 | 0/3 | 0/3 | 0/3 | 0/1 | 0/2 |
| CC H7-8 | ↗ | 0/1 | 0/1 | 0/8 | 0/8 | 0/8 | 0/8 | 0/6 | 0/6 |
| | = | 1/1 | 1/1 | 6/8 | 8/8 | 5/8 | 8/8 | 3/6 | 6/6 |
| | ↘ | 0/1 | 0/1 | 2/8 (8.08%) | 0/8 | 3/8 (6.13%) | 0/8 | 3/6 (6.71%) | 0/6 |
| SCR H7-8 | ↗ | 0/2 | 2/2 (43.58%) | 0/8 | 5/8 (98.68%) | 0/7 | 3/7 (19.04%) | 0/7 | 0/7 |
| | = | 0/2 | 0/2 | 3/8 | 3/8 | 4/7 | 4/7 | 2/7 | 7/7 |
| | ↘ | 2/2 (12.47%) | 0/2 | 5/8 (9.58%) | 0/8 | 3/7 (8.70%) | 0/7 | 5/7 (10.55%) | 0/7 |

| Right limbs Leading | | LLG | | LIP | | LBFP | |
|---|---|---|---|---|---|---|---|
| | | Onset | Amplitude | Onset | Amplitude | Onset | Amplitude |
| CC INTACT | ↗ | 0/7 | 0/6 | 3/3 (5.35%) | 1/3 (24.24%) | 0/8 | 5/8 (3.09%) |
| | = | 6/7 | 5/6 | 0/3 | 2/3 | 3/8 | 3/8 |
| | ↘ | 1/7 (4.33%) | 1/6 (14.85%) | 0/3 | 0/3 | 5/8 (4.16%) | 0/8 |
| CC H1-2 | ↗ | 0/6 | 0/6 | 2/2 (8.88%) | 0/2 | 0/5 | 0/5 |
| | = | 6/6 | 6/6 | 0/2 | 1/2 | 5/5 | 5/5 |
| | ↘ | 0/6 | 0/6 | 0/2 | 1/2 (32.03%) | 0/5 | 0/5 |
| SCR H1-2 | ↗ | 0/7 | 0/6 | 1/3 (13.84%) | 1/3 (31.55%) | 0/7 | 4/7 (15.31%) |
| | = | 3/7 | 5/6 | 2/3 | 2/3 | 7/7 | 3/7 |
| | ↘ | 4/7 (8.83%) | 1/6 (36.99%) | 0/3 | 0/3 | 0/7 | 0/7 |
| Other H1-2 | ↗ | 0/5 | 0/5 | 0/1 | 0/1 | 0/2 | 0/2 |
| | = | 5/5 | 5/5 | 1/1 | 1/1 | 2/2 | 2/2 |
| | ↘ | 0/5 | 0/5 | 0/1 | 0/1 | 0/2 | 0/2 |
| CC H7-8 | ↗ | 0/6 | 0/6 | 0/2 | 0/2 | 0/7 | 3/7 (12.95%) |
| | = | 6/6 | 6/6 | 2/2 | 2/2 | 7/7 | 4/7 |
| | ↘ | 0/6 | 0/6 | 0/2 | 0/2 | 0/7 | 0/7 |
| SCR H7-8 | ↗ | 0/7 | 0/7 | 1/3 (9.32%) | 0/3 | 0/7 | 4/7 (14.02%) |
| | = | 4/7 | 5/7 | 2/3 | 2/3 | 6/7 | 3/7 |
| | ↘ | 3/7 (7.25%) | 2/7 (22.33%) | 0/3 | 1/3 (17.41%) | 1/7 (7.88%) | 0/7 |

**Table S2. Modulation of EMG burst onset and amplitude before and after lateral hemisection during obstacle negotiation with Right limbs leading.** The table shows the number of individual cats/total number of cats with EMGs available for a given muscle. Upward and downward arrows indicate significant increases or decreases, respectively, while the equal sign represents no significant change (two-factor repeated measures ANOVA). The percent value indicates the average of all cats that showed a significant increase or decrease.

| | | Time | Height | Interaction |
|---|---|---|---|---|
| Type of negotiation | CC | <0.0001 | <0.0001 | <0.0001 |
| | SCR | 0.0329 | <0.0001 | |
| | Other | <0.0001 | 0.4972 | |
| | | **Side** | **Height** | **Interaction** |
| Side CC | Intact | 0.3413 | 0.7694 | |
| | H1-2 | 0.0054 | <0.0001 | 0.0112 |
| | H7-8 | <0.0001 | 0.3811 | |
| Side SCR | H1-2 | 0.5414 | 0.9999 | |
| | H7-8 | 0.6607 | 0.9999 | |
| Side Other | H1-2 | 0.0069 | 0.9999 | |
| | H7-8 | 0.4836 | 0.9999 | |

**S3 Table. P values for each negotiation type and side preference.** P values comparing time and heights are indicated (mixed-effects model ANOVA) for each negotiation type (Fig. 1B). P values comparing side and heights are indicated (mixed-effects model ANOVA) for each type of negotiation (Fig. 1C).

| | Left limbs leading | | Right limbs leading | |
|---|---|---|---|---|
| | Negotiation | Height | Negotiation | Height |
| Approach distance | <0.0001 | 0.0135 | 0.3345 | 0.2827 |
| Reception distance | 0.0002 | 0.3272 | 0.0031 | 0.3467 |
| Difference of clearance | <0.0001 | 0.0151 | <0.0001 | 0.2397 |

**S4 Table. P values for distances from the obstacles and difference of clearance.** P values comparing each negotiation type and heights after hemisection to complete clearance in intact cats are indicated (mixed-effects model ANOVA) for approach distance, reception distance and difference of clearance (Fig. 3).

| | | Left limbs leading | | Right limbs leading | |
|---|---|---|---|---|---|
| | | Step | Height | Step | Height |
| Hip angular excursion | CC Intact | 0.7717 | 0.6414 | 0.0927 | 0.8841 |
| | CC H1-2 | 0.6006 | 0.6980 | <0.0001 | 0.5320 |
| | SCR H1-2 | 0.0151 | 0.2633 | 0.0317 | 0.9682 |
| | Other H1-2 | 0.0428 | 0.3388 | 0.0477 | 0.7446 |
| | CC H7-8 | 0.0848 | 0.6634 | 0.2055 | 0.4734 |
| | SCR H7-8 | 0.4621 | 0.6200 | 0.5813 | 0.3305 |
| Knee angular excursion | CC Intact | <0.0001 | 0.0104 | <0.0001 | 0.0112 |
| | CC H1-2 | <0.0001 | 0.1908 | 0.0024 | 0.8475 |
| | SCR H1-2 | 0.0003 | 0.1699 | 0.0007 | 0.0335 |
| | Other H1-2 | 0.0001 | 0.2059 | 0.0005 | 0.1037 |
| | CC H7-8 | <0.0001 | 0.0395 | 0.0038 | 0.0202 |
| | SCR H7-8 | <0.0001 | 0.0379 | <0.0001 | 0.0776 |
| Ankle angular excursion | CC Intact | <0.0001 | 0.0164 | <0.0001 | 0.0129 |
| | CC H1-2 | 0.0007 | 0.6119 | 0.3352 | 0.5250 |
| | SCR H1-2 | 0.0005 | 0.5733 | 0.0010 | 0.5052 |
| | Other H1-2 | 0.0002 | 0.1432 | 0.0995 | 0.6648 |
| | CC H7-8 | <0.0001 | 0.6309 | 0.7376 | 0.0137 |
| | SCR H7-8 | 0.0046 | 0.1939 | 0.0001 | 0.7783 |

**S5 Table. P values for angle excursions.** P values comparing steps and heights are indicated (mixed-effects model ANOVA) for each type of negotiation for hip, knee and ankle angular excursions (Fig. 5).

| | Left limbs leading | | Right limbs leading | |
|---|---|---|---|---|
| | **Negotiation** | **Height** | **Negotiation** | **Height** |
| **Hip** | 0.0005 | 0.9726 | <0.0001 | 0.8704 |
| **Knee** | 0.0357 | 0.8675 | <0.0001 | 0.4480 |
| **Ankle** | 0.0026 | 0.3240 | <0.0001 | 0.0318 |

**S6 Table. P values for timing of joint flexions.** P values comparing each negotiation type and obstacle heights to complete clearance in intact cats are indicated (mixed-effects model ANOVA) for the timing of hip, knee and ankle joint flexion (Fig. 6).

| | Left limbs leading | | Right limbs leading | |
|---|---|---|---|---|
| | **Negotiation** | **Height** | **Negotiation** | **Height** |
| **Period 1** | <0.0001 | 0.8622 | <0.0001 | 0.7857 |
| **Period 2** | <0.0001 | 0.6679 | <0.0001 | 0.7149 |
| **Period 3** | 0.0046 | 0.8874 | 0.6917 | 0.3030 |
| **Period 4** | <0.0001 | 0.8979 | 0.0007 | 0.1006 |
| **Period 5** | 0.0002 | 0.3649 | 0.8811 | 0.4124 |
| **Period 6** | <0.0001 | 0.3728 | <0.0001 | 0.7671 |
| **Period 7** | 0.4226 | 0.8362 | 0.5083 | 0.7347 |
| **Period 8** | <0.0001 | 0.7496 | <0.0001 | 0.3322 |
| **Period 9** | 0.5029 | 0.6289 | 0.0024 | 0.7250 |

**S7 Table. P values for support periods.** P values comparing each negotiation type and obstacle heights to complete clearance in intact cats are indicated (mixed-effects model ANOVA) for support periods (Fig. 7).

| | | Left limbs leading | | Right limbs leading | |
|---|---|---|---|---|---|
| | | **Step** | **Height** | **Step** | **Height** |
| **Speed** | **CC Intact** | <0.0001 | 0.7100 | <0.0001 | 0.7247 |
| | **CC H1-2** | 0.7912 | 0.1926 | 0.0532 | 0.5841 |
| | **SCR H1-2** | 0.3623 | 0.8206 | 0.9531 | 0.1982 |
| | **Other H1-2** | 0.5278 | 0.0608 | 0.0557 | 0.3878 |
| | **CC H7-8** | 0.8201 | 0.0595 | 0.2215 | 0.1364 |
| | **SCR H7-8** | 0.4673 | 0.5434 | 0.1487 | 0.8451 |
| **Stride length** | **CC Intact** | <0.0001 | 0.8624 | <0.0001 | 0.8081 |
| | **CC H1-2** | 0.0445 | 0.6628 | <0.0001 | 0.5082 |
| | **SCR H1-2** | 0.0322 | 0.3638 | 0.0001 | 0.8381 |
| | **Other H1-2** | 0.0526 | 0.2662 | 0.0026 | 0.9026 |
| | **CC H7-8** | <0.0001 | 0.3722 | 0.0472 | 0.2348 |
| | **SCR H7-8** | 0.4853 | 0.8083 | 0.0006 | 0.8864 |

**S8 Table. P values for speed and stride length.** P values comparing the control and obstacle steps and heights are indicated (mixed-effects model ANOVA) for each type of negotiation for speed and stride length (Fig. 8).
